# Context-dependent decision making in a premotor circuit

**DOI:** 10.1101/757104

**Authors:** Zheng Wu, Ashok Litwin-Kumar, Philip Shamash, Alexei Taylor, Richard Axel, Michael N. Shadlen

## Abstract

Cognitive capacities afford contingent associations between sensory information and behavioral responses. We studied this problem using an olfactory delayed match to sample task whereby a sample odor specifies the association between a subsequent test odor and rewarding action. Multi-neuron recordings revealed representations of the sample and test odors in olfactory sensory and association cortex, which were sufficient to identify the test odor as match/non-match. Yet, inactivation of a downstream premotor area (ALM), but not orbitofrontal cortex, confined to the epoch preceding the test odor, led to gross impairment. Olfactory decisions that were not context dependent were unimpaired. Therefore, ALM may not receive the outcome of a match/non-match decision from upstream areas but contextual information—the identity of the sample—to establish the mapping between test odor and action. A novel population of pyramidal neurons in ALM layer 2 may mediate this process.

## Introduction

Brain functions deemed “cognitive” exhibit complexities that extend an organism’s behavioral repertoire beyond simple sensory-response associations, motor programs, and instructed actions. Cognitive functions exploit contingent, hierarchical processes of decision making and executive control. For example, instead of choosing an action, a decision might lead an organism to choose a strategy, or to switch tasks, or to make yet another decision. Cognitive functions may also transpire over flexible time scales without precipitating an immediate behavior, as when a decision is based on information acquired at some moment and combined with information acquired later.

Neural mechanisms of decision making and executive control have been studied primarily in the parietal and prefrontal cortex of nonhuman primates. These studies focused mainly on neurons that exhibit responses over long time-scales and implicated processes such as working memory (Funahashi et al., 1989; Fuster and Alexander, 1971), planned action (Cisek and Kalaska, 2005; Evarts and Tanji, 1976; Snyder et al., 1997), behavioral state (Harvey et al., 2012; Kadohisa et al., 2013), representation of stimulus qualities (Freedman and Assad, 2006; Romo et al., 1999), and reasoning (Gold and Shadlen, 2007; Yang and Shadlen, 2007). Thus the study of the neural mechanisms of perceptual decision making may provide an initial logic for higher order cognition. This will require an understanding of neural circuits at a level that is not yet possible to achieve in nonhuman primates. Thus there has been growing interest in the pursuit of elementary cognitive functions in the mouse, for which genetic and viral tools for circuit manipulation abound (Carandini and Churchland, 2013; Luo et al., 2018). The challenge is to find simple behaviors, within the mouse repertoire, that have the potential to elucidate more complex cognitive functions.

We developed a simple task that allows us to study rudiments of executive control and decision-making in a mouse. The task is a variant of the logical XOR problem, realized as an olfactory delayed match to sample task (DMS; Fig. 1) (Liu et al., 2014). The mouse is exposed to a *sample odor*, either S_A_ or S_B_, and, after a short delay, receives either A or B as a test order. To receive a reward, the mouse must lick to the left if the sample and test odors are the same and to the right if they are different. We reasoned that mice could perform this task by interpreting the **sample odor** as an instruction. The instruction is to select the appropriate association between the **test odor** and the correct behavioral response—a left or right lick.

**Figure 1:**
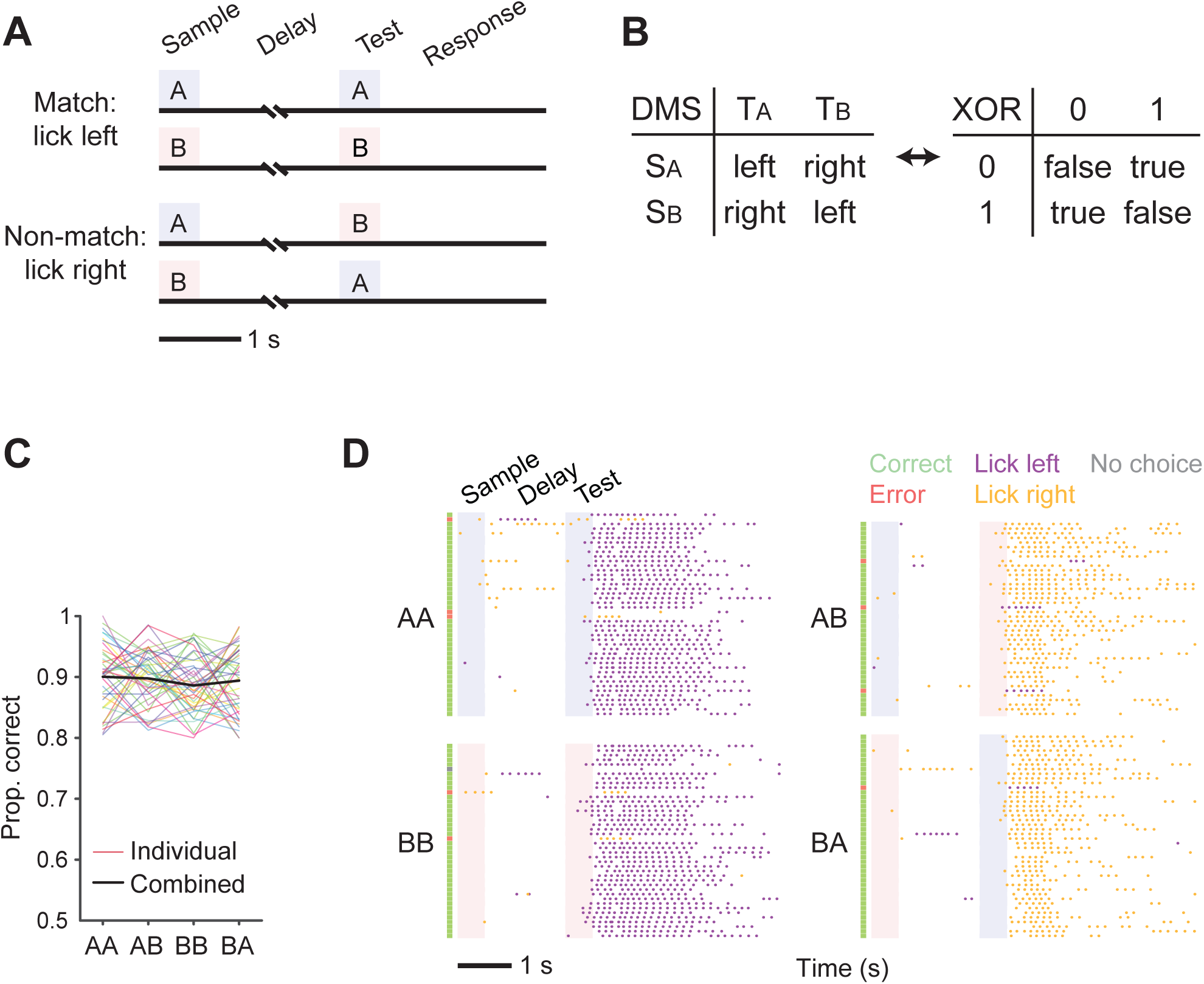
Lick-left/lick-right olfactory delayed match to sample task. (**A**) Task structure. Mice are presented with two odorants (A: pinene, B: hexenol) separated by a delay and must decide whether they are the same (match) or different (non-match). The two odors create four unique pairs, or trial types. Match trials are rewarded on the left port and non-match on the right (see Methods). (**B**) The DMS task is a variant of the logical XOR problem. The outcome/response depends on whether the two bits of information are the same or not. (**C**) The proportion of correct trials in well-trained animals. Each colored line represents data from one animal and the black line is the mean of all animals (n = 41). (**D**) An example behavior session from a well-trained animal. The four trial types are randomly presented in the session but grouped here for plotting. Each row is a trial. The green, red, and gray markers on the left side of each trial denote respectively the outcome of correct, error, and no choice. The A and B odors are indicated by the light blue and light red shadings. The magenta and yellow tick marks are the left and right licks, respectively. Mice were trained to suppress their premature licks during the sample and delay epochs. The sample and test epochs are 0.5 s each and the delay is 1.5 s. See also Figure S1.

We hypothesized that this hierarchical control might be solved by changing the configuration of cortical circuitry in the premotor cortex, area ALM. Studies from Svoboda and colleagues have shown that ALM plays an essential role in behaviors in which a sensory cue serves as an instruction to lick to the left or right (Guo et al., 2014; Li et al., 2015; Svoboda and Li, 2018). They showed that many neurons in ALM can hold such an instruction—or the plan to lick to the left or right— in persistent activity through a delay period. In the DMS task we study, the sample odor cannot provide an instruction to lick left or right, but only a context to interpret the test odor. Thus the mouse cannot plan a lick until the arrival of the test odor after the delay. We investigated three hypotheses for the involvement of ALM in this task. First, areas upstream to ALM could solve the match/non-match discrimination and project this solution to ALM which organizes an appropriate lick response. Second, ALM could decode upstream representations that combine sample and test odors to select the appropriate response. Third, ALM could receive information about identity of the sample odor to instantiate the appropriate response to the test odor. We provide experimental evidence for this third possibility. We show that the sample odor is represented in ALM itself, and that this representation allows ALM to associate the test odor with the appropriate lick response.

## Results

Mice were trained to compare a sample and test odor, separated by a delay and to report their decisions of “match” or “non-match” by licking to the left or to the right, respectively (Fig. 1A). The lick-left, lick-right design requires distinct actions to report both match and non-match. Importantly, animals cannot solve the task until they smell the test odor. For most experiments, we used the same two odors, (+)-α-Pinene and cis-3-Hexen-1-ol (odors A and B, respectively) on all trials. Animals were trained to suppress their premature licks before test onset (Fig. 1D) and do not appear to use licks to represent and remember the sample odors (Fig. S1E-G).

Mice performed the task with a median accuracy of 90% (n = 41; Fig. 1C). When challenged with a novel pair of odorants after training with pinene and hexenol, mice performed at chance level with a high no-choice rate (Fig. S1C). This suggests that the mice failed to generalize the match/non-match rule. Rather they learned that *test odor T_A_* instructs “lick left” if preceded by *sample odor S_A_* (AA trial), whereas it instructs “lick right” if preceded by *sample odor S_B_* (BA trial). The complementary logic holds for the *test odor T_B_* (Fig. 1B). This flexible association between test odor and licking response is contingent on the sample odor identity, which must be represented through the delay in order to affect the match/non-match decision.

We first characterize the neural representation of the sample and test odors by surveying neural responses in the Piriform cortex (Pir), the orbitofrontal cortex (OFC) and the anterolateral motor cortex (ALM) while the mouse performed the DMS task. We wished to determine whether each of these areas contain a persistent representation of the sample odor and whether they encode the test odor in a way that might inform the match-nonmatch decision. As detailed in the next section, the findings support the possibility that match/non-match decision can be established within Pir and OFC and then transmitted to the ALM, to render the lick response. We then test this model by inactivating ALM during the sample and delay periods of the task. We find evidence against this model and instead establish a necessary role of ALM during the sample and delay periods. Finally, using 2-photon Ca imaging, we expose a class of neurons in superficial ALM that could mediate the circuit changes required to allow ALM to associate the test odor with an appropriate lick response.

### Representations of sensory and motor signals in Pir, OFC and ALM

We performed electrophysiological recordings in brain areas likely to be involved in the DMS task, piriform cortex (Pir), orbitofrontal cortex (OFC), and ALM. Pir is a primary olfactory sensory area (Giessel and Datta, 2014; Sosulski et al., 2011) that encodes odor identity. Pir projects to OFC, a higher-order associative area that is thought to encode value, expectation, and working memory (Bechara et al., 2000; Mainen and Kepecs, 2009; Padoa-Schioppa and Assad, 2006; Ramus and Eichenbaum, 2000). Figure 2 provides examples of neurons that respond selectively to (*i*) one or the other sample odor during the sample- and delay-periods (1^st^ and 2^nd^ columns); (*ii*) one or the other test odor (3^rd^ column); and (*iii*) one or the other choice (4^th^ column). It also shows the strength of the selectivity across all the neurons recorded. The selectivity index quantifies the degree to which the distributions of firing rates to the two conditions (e.g., odor A or B) are non-overlapping (see Methods). In what follows we refer to a neuron as selective if the difference in the mean responses is statistically reliable (Mann-Whitney U test, p<0.01, two-tailed, uncorrected; indicated by shading in Fig. 2D-G).

**Figure 2:**
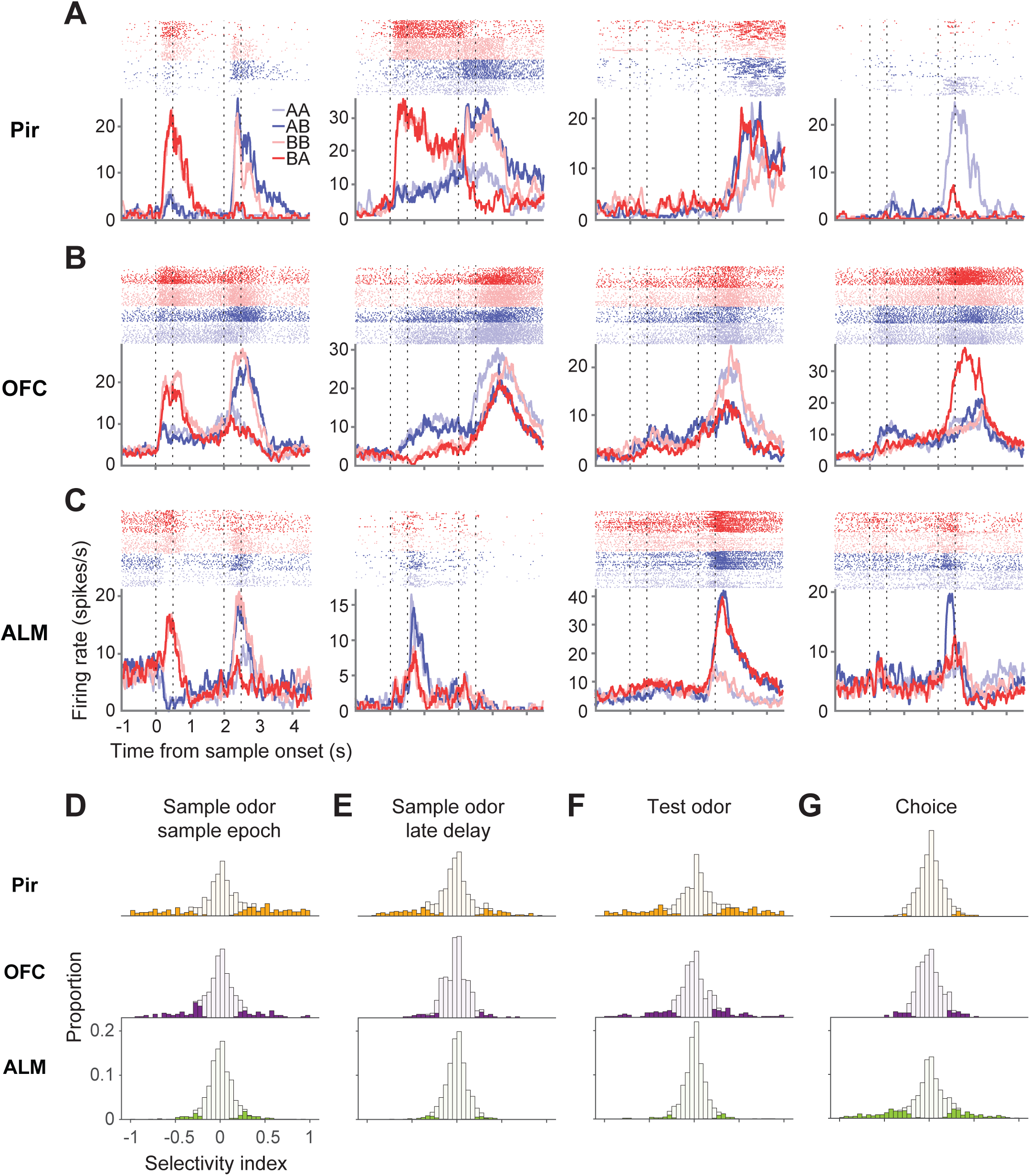
Neural recordings of Pir, OFC and ALM. (**A-C**) Firing rates of example neurons from Pir (A), OFC (B), and ALM (C) recordings. The rasters (top) show the times of action potentials in individual trials (rows). Traces are mean firing rates (100 ms bins) across trials. The dotted lines indicate the sample and test epochs hereafter. (**D-G**) Selectivity indices of Pir, OFC, and ALM neurons for sample odor during the sample epoch (D), sample odor during the last 0.5 s of the delay epoch (E), test odor during the test epoch (F), and choice during the test epoch (G). n = 647, 380 and 1086 neurons for Pir, OFC and ALM, respectively. Shading indicates neurons that are selective for an odor (D-F) or behavioral choice (G) determined by Mann-Whitney U test, *P* < 0.01, not corrected for multiple comparisons. See also Figure S2.

More than a third of neurons (37%) in Pir responded selectively to either odor A or odor B during the sample epoch (Fig. 2A, D). Of these, 71% exhibited the same odor preference during the sample and test epochs (Fig. S2A) suggesting that these neurons represent sensory information and encode odor identity. Most of the sample-selective neurons responded transiently (e.g., Fig. 2A, left most panel), but 28% exhibited persistent selectivity through the delay (e.g., Fig. 2A, 2^nd^ panel from left; Fig. S2D). These neurons can therefore inform downstream neurons of the identity of the sample odor at the time of test. A subset of Pir neurons responded to the test odor in a way that depended on the identity of the sample odor (Fig. S2G; Methods). These neurons are characterized as trial type selective (6.2% of all Pir neurons). Finally, 4.5% of Pir neurons exhibited selective responses during the test epoch that reflect either a match or non-match between sample and test odors (Fig. 2A, G). The information represented by these last two classes of neurons, trial type and match/non-match, would appear to be sufficient to guide motor output in downstream brain areas.

In OFC, 24.5% of the neurons were odor selective during the sample epoch (Fig. 2B, D) and 60% of these exhibited the same odor preference at test (Fig. S2B), suggesting that they encode odor identity. Compared to Pir, fewer of the sample-selective neurons in OFC exhibited persistent spiking through the delay epoch (Fig. S2E; 13% in OFC versus 28% in Pir). Only 3.7% of the neurons were selective to trial type (Fig. S2H), but a greater fraction (11%) responded selectively to match/non-match trials (Fig. 2G). Therefore, the OFC also appears to have integrated sensory inputs in a way that could guide appropriate motor output.

Unlike Pir and OFC, which largely represent odor identity, the dominant task related activity in ALM was correlated with the lick response—the outcome of the match/non-match decision; 28% of ALM neurons distinguished match from non-match trials during the test epoch (Fig. 2C, G). Analyses of error trials and premature licks show that the neural activity reflects lick direction rather than the identities of the sample and test odors or the true match/non-match condition, consistent with the role of ALM in motor planning (Fig. S2J, K) (Guo et al., 2014; Li et al., 2015; Svoboda and Li, 2018). A largely non-overlapping population in ALM (12%) exhibited weak odor selectivity during the sample epoch, and among these, 12% (less than 1.5% overall) maintained this selectivity through the end of the delay epoch (Fig. 2D, S2F). Very few neurons exhibited a selective response to the test odor, and only 1.4% of ALM neurons were trial type selective (Fig. S2I). Together, these recordings are consistent with the known role for ALM in the preparation of an appropriate motor response (Svoboda and Li, 2018) following the match/non-match decision.

This survey of a sensory, association and premotor cortex reveals neurons in each area that exhibit one or more of the properties required to solve the DMS task: (*i*) a representation of the sample odor during the delay epoch and (*ii*) a response to the test odor that is possibly modulated by the identity of the sample odor and (*iii*) the conversion of these sensory signals to choice-related activity. Figure 3A shows the averaged difference in firing rate to each neuron’s preferred versus nonpreferred odor during the trial. The assignment of preferred odor was derived from five randomly selected trials to each odor, which are excluded from the averages. The procedure ensured an unbiased estimate of the difference (see Methods). The average difference is strongest in Pir and weakest in ALM. In the left panel, the magnitude of the difference reflects a combination of greater selectivity and the fraction of neurons that are selective, as nonselective neurons drive the average toward zero. In the right panel, the averages comprise only the selective cases from each area (filled histograms in Fig. 2). All three areas contain signals that could convey the representation of the sample odor (Pir > OFC > ALM). However, this conclusion is based on averaged activity across trials.

**Figure 3:**
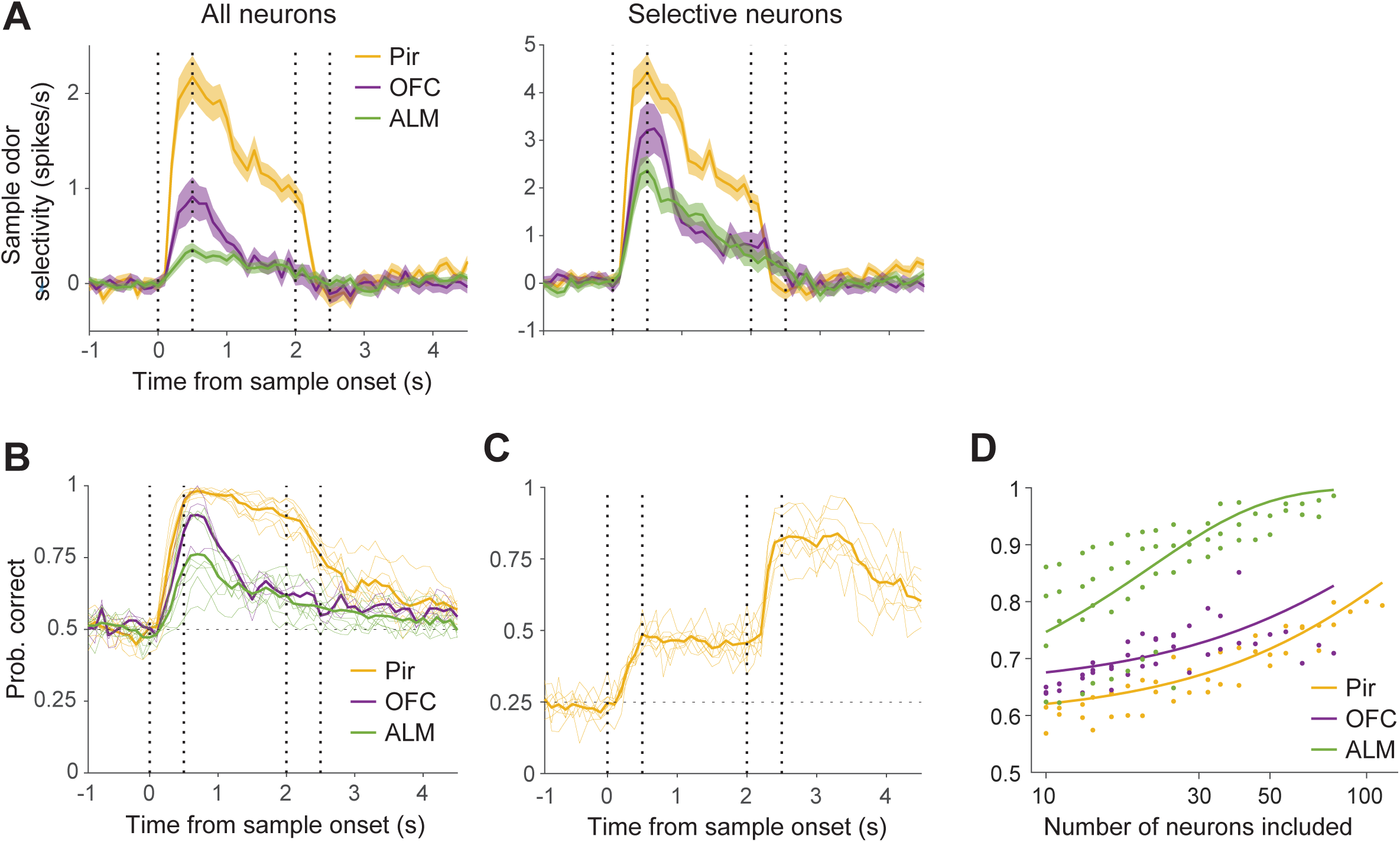
Odor selectivity and decoding task-relevant features using the neural responses. (**A**) Population sample odor selectivity of Pir, OFC, and ALM neurons. The ordinate is the difference in firing rate to the preferred and the nonpreferred odor (Methods). The lines represent the means across neurons and the shading represents the s.e.m. *Left graph*, all recorded neurons from each area contribute to the averages. *Right graph*, only neurons that are selective for sample odor in the sample and delay epochs contribute to the averages (n = 319, 105 and 147 neurons for Pir, OFC and ALM, respectively). (**B**) Performance of a support vector machine trained to classify the sample odor identity using simultaneously recorded neurons from each of the three areas. Each thin line represents the probability of correct classifications of the “held out” trials using data from one session (see Methods). Thick lines represent the mean of all sessions from an area (color). (**C**) Performance of a classifier of trial type using all simultaneously recorded Pir neurons from each session. Same conventions as in (A). (**D**) Performance of a classifier of match/non-match trials using the neural responses after test odor onset and before animal’s first lick. Each point represents the performance using *N* randomly selected neurons as input to the classifier, calculated using sessions in which at least *N* neurons were recorded simultaneously. The traces are fits to the classifier performance (logistic regression). For each area, the performance improves as more neurons are included.

To determine the information available about task variables present in population responses, we trained linear classifiers to decode these variables from simultaneously recorded neurons on single trials (from 14 to 120 simultaneously recorded neurons; see Methods). As shown in Figure 3B, all three areas contain signals that support classification of the sample odor above chance levels throughout the sample and delay periods. The performance of the classifier is lower for ALM recordings compared to Pir recordings, for which the sample odor identity can be decoded throughout the delay period and into the test epoch. This extended sample-selective response could potentially be used to perform a comparison between sample and test odors. Consistent with this, classification of all four trial types is possible using the Pir recordings (Fig. 3C). In this same epoch (test odor onset to the first lick response), it is possible to decode the binary choice from all three areas, with ALM exhibiting the highest performance (Fig. 3D).

The neural recordings and decoding analyses would appear to support a traditional hierarchical view of the mechanism responsible for resolving the match/non-match decision. That is, the expression of the decision by the left- and right-lick neurons in ALM could arise by reading out activity in OFC or Pir, perhaps via another intermediate area between Pir and ALM. On this view, area ALM is only essential once the test odor arrives, when it must either (*i*) convert the decision to a lick response or (*ii*) perform a computation, similar to our decoder, to convert the population response from upstream areas to a left or right lick response. Both mechanisms lead to the prediction that inactivation of ALM during the sample and delay periods should not impair performance on the task. As we next show, this seemingly obvious prediction is incorrect.

### ALM is required for the match/non-match computation

We inactivated ALM bilaterally on a fraction of the trials by photostimulation of GABAergic interneurons expressing channelrhodopsin-2 (ChR2) (Fig. 4A, S3; Methods) (Guo et al., 2014; Zhao et al., 2011). Inactivation was confined to the sample and delay epochs and was tapered gradually at the end of the delay (see Methods) such that by the onset of the test odor, ALM should be capable of receiving information from upstream sensory and association areas. We reasoned that if ALM reads out the DMS decision computed in upstream areas then inactivation of ALM before the arrival of the test odor should not impair performance.

**Figure 4:**
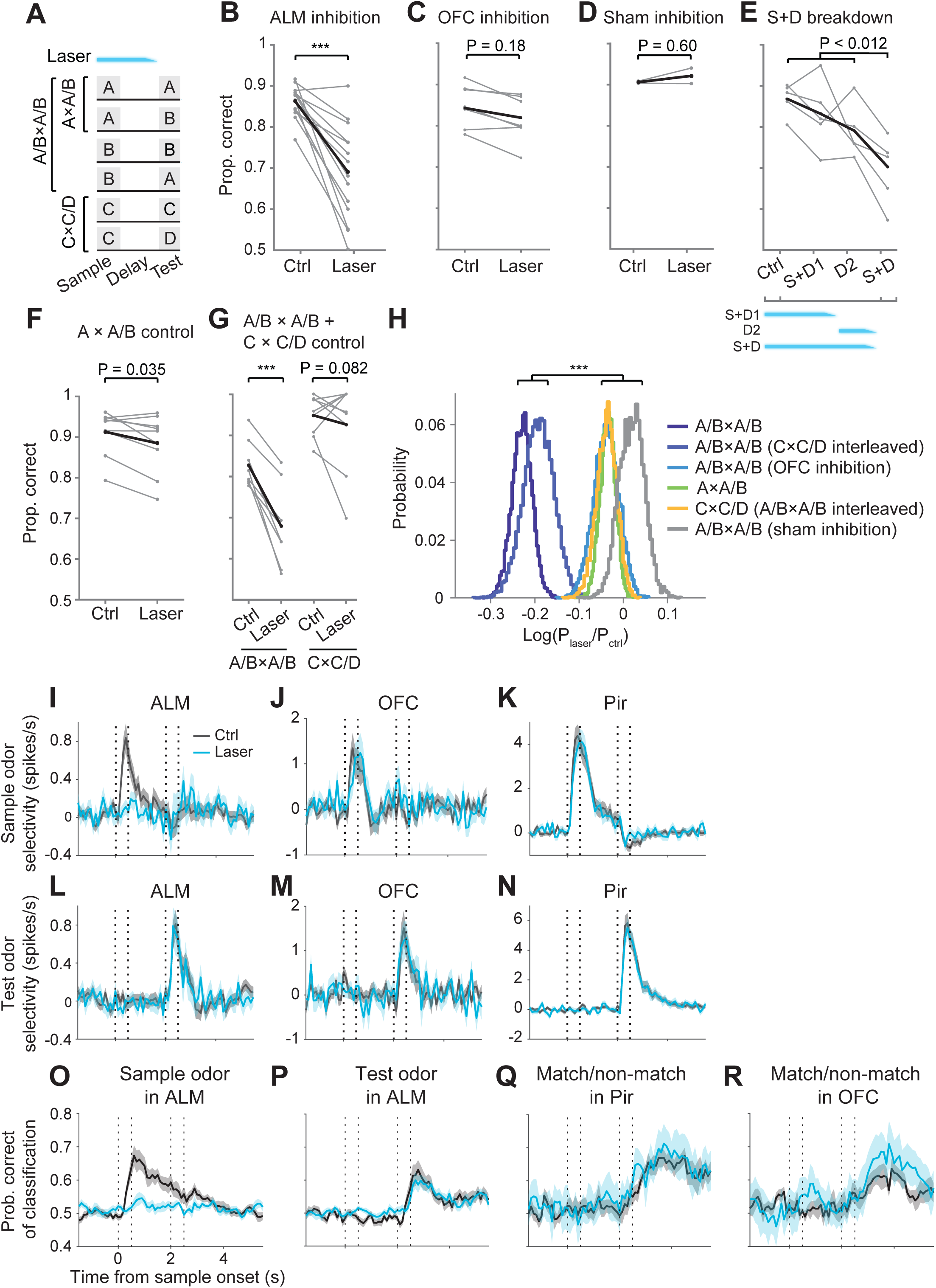
Optogenetic inactivation experiments. (**A**) Schematic of the DMS and control tasks employed in the inhibition experiments (time not drawn to scale). A/B × A/B design is the same DMS task as in the recording experiments. On inactivation trials, unless otherwise indicated, the laser was on for 2 s including a 0.25 s power ramp-down at the end (tapered blue bar). In the interleaved A/B × A/B & C × C/D task, the delay epoch is 4 s (Methods). (**B**) Proportion of correct trials in the A/B × A/B task with and without bilateral ALM inactivation. Each thin line represents data from one animal. The thick line is the combined performance from 4092 trials (14 animals). Statistical reliability of effects in panels (B-G) are based on permutation tests (*, ** and *** denote *P* < 0.01, 0.001 and 0.0001). (**C**) Proportion of correct trials in the A/B × A/B task with and without bilateral OFC inactivation (7 animals, 2278 trials). (**D**) Proportion of correct trials in the A/B × A/B task by ChR2 non-carriers with and without sham ALM inactivation (2 animals, 437 trials). (**E**) Effect of inactivation during portions of the sample and delay epochs (5 animals, 3732 trials). Same conventions as in (B-D). *Ctrl*, no inactivation; *S+D1*, sample and first 1.5 s of the delay; *D2*, last 1.0 s of the delay; *S+D*, sample and delay. (**F**) Proportion of correct trials in the A × A/B control task with and without bilateral ALM bilateral inactivation (9 animals, 2198 trials). (**G**) Proportion of correct trials in the interleaved A/B × A/B & C × C/D task with and without bilateral ALM inactivation (9 animals, 2728 trials). (**H**) ALM inactivation caused greater impairment on the A/B × A/B task than on the control tasks and OFC inactivation on the same A/B × A/B task. The effect of inactivation is expressed as the log odds (LO) of the mouse performance with and without inactivation (Methods, Eq. 4). Distributions of the LO were estimated by a bootstrap procedure. The largest effects were evident when ALM was inactivated and the sample was informative (*t*-test, *d.f.* determined by size of data set). (**I-K**) Population sample odor selectivity of ALM, OFC, and Pir with and without ALM inactivation. The ordinate is the difference in firing rate to the preferred and the nonpreferred odor across neurons (see Methods). Note the different scales. Shading is s.e.m. n = 470, 104 and 188 neurons for ALM, OFC and Pir, respectively. The same neurons are used in the analyses below. (**L-N**), Population test odor selectivity of ALM, OFC, and Pir with and without ALM inactivation. (**O-R**) Performance of the SVM decoder to classify sample odor identity using ALM responses (O), test odor identity using ALM responses (P), match/non-match using Pir response (Q), and match/non-match using OFC responses (R) with and without ALM inactivation. See also Figure S3 and S4.

Contrary to predictions, bilateral inactivation of ALM markedly impaired performance on the DMS task (Fig. 4B). The proportion of correct trials decreased from 0.86 on the control trials to 0.68 (0.5 reflects chance performance). In contrast, bilateral inactivation of OFC induced no impairment on the same task (proportion correct: 0.82 and 0.84 with and without inactivation; Fig. 4C). Photostimulation of ALM in animals that did not express ChR2 did not diminish performance (Fig. 4D). Inactivation of ALM caused less impairment when it was restricted to only a portion of the sample plus delay epochs (Fig. 4E). Inactivation during the late delay reduced the proportion of correct trials from 0.86 to 0.79, whereas inactivation during the sample and early delay reduced the proportion to 0.83. These more modest effects suggest that the impairment can be ameliorated when there is a time window in which ALM can receive task related information.

The behavioral impairment induced by ALM inactivation is a consequence of altered activity within ALM itself during the sample and delay. Photoinactivation of ALM did not affect odor or match/non-match selectivity in upstream areas, Pir and OFC. During inactivation, the population odor selectivity in Pir or OFC did not change appreciably (Fig. 4J, K, M, N) and our ability to decode match vs non-match from activity was similarly unchanged (Fig. 4Q, R). Neural recordings from ALM during silencing demonstrated that optogenetic suppression eliminated the representation of the sample odor, but neural activity returns after the laser was ramped down (Fig. 4I, L, O, P). Despite their diminished performance, animals invariably licked to one of the two ports in inactivation trials, and there was no consistent change in the reaction time (Fig. S4A, B). From these observations, we conclude that the impairment was not explained by a loss of processing capacity in Pir or OFC or from physiological sequelae of photoinactivation of ALM that might affect its function during the test epoch. This last conclusion deserves further scrutiny.

A possible concern is that the inactivation of ALM during sample and delay epochs disrupts the ALM circuitry such that it is unable to process information about the test odor or to receive information about the decision from an upstream area. We evaluated this possibility with two control experiments that required the mice to lick to the left or right, based on the identity of a test odor, but did not require a comparison of the test odor to the sample odor. In the first control, an uninformative sample odor A was presented on all trials, but the mouse was rewarded for licking left or right for test odor A or B, respectively (AA and AB trials; Fig. 4A). Inactivation during the sample and delay epochs produced a small impairment (0.91 to 0.88 correct; Fig. 4F) that was much less pronounced than on the DMS trials (Fig. 4H, green vs. dark blue, p<0.0001; Methods).

In the second control, we incorporated two additional trial types into the AB × AB design using two new odors (C and D). In these CC and CD trials, as in the previous control, the correct lick behavior was determined only by the test odor. The six trial types were randomly interleaved (Fig. 4A). Inactivation of ALM during the sample and delay epochs led to minimal impairment in CC and CD trials (0.94 to 0.91 correct), whereas significant impairment was replicated in the interleaved DMS trials (0.82 to 0.68 correct; Fig. 4G). Further, the laser introduced no side bias in the CC/CD control (*β_2_* = −0.46, s.e.m. = 0.40, *P* = 0.25, Equation 8; see Methods), demonstrating that the bias observed in some of the DMS trials is not due to unbalanced inactivation of the two hemispheres (see also Fig. S4C). These control experiments demonstrate that after recovery from inactivation, ALM is capable of processing information that instructs a licking response via a simple association between two odors and two actions. The impairment on the DMS task must therefore arise by interfering with the process that allows the sample odor to establish the association on each trial. Moreover, it implies that this process occurs in ALM.

### Enrichment of sample-selective neurons in ALM Layer 2

Based on the neural recordings, the necessity of ALM during the sample and delay seems highly perplexing. Less than 1.5% of neurons in ALM had activity that was informative about the identity of the sample. We considered that we might have missed neurons, especially from superficial cortical layers (Fig. S2L; see also Guo et al., 2017, their Fig. 3D). We therefore examined neural responses in ALM with 2-photon calcium imaging while mice performed the DMS task. Imaging was performed in mice expressing the calcium indicator GCaMP6f in pyramidal cells (Chen et al., 2013; Madisen et al., 2015) (Methods). Consistent with our electrical recordings, calcium imaging revealed that ∼37% of the neurons were choice selective across all cortical depths examined (Fig. 5G, S5A, D).

**Figure 5:**
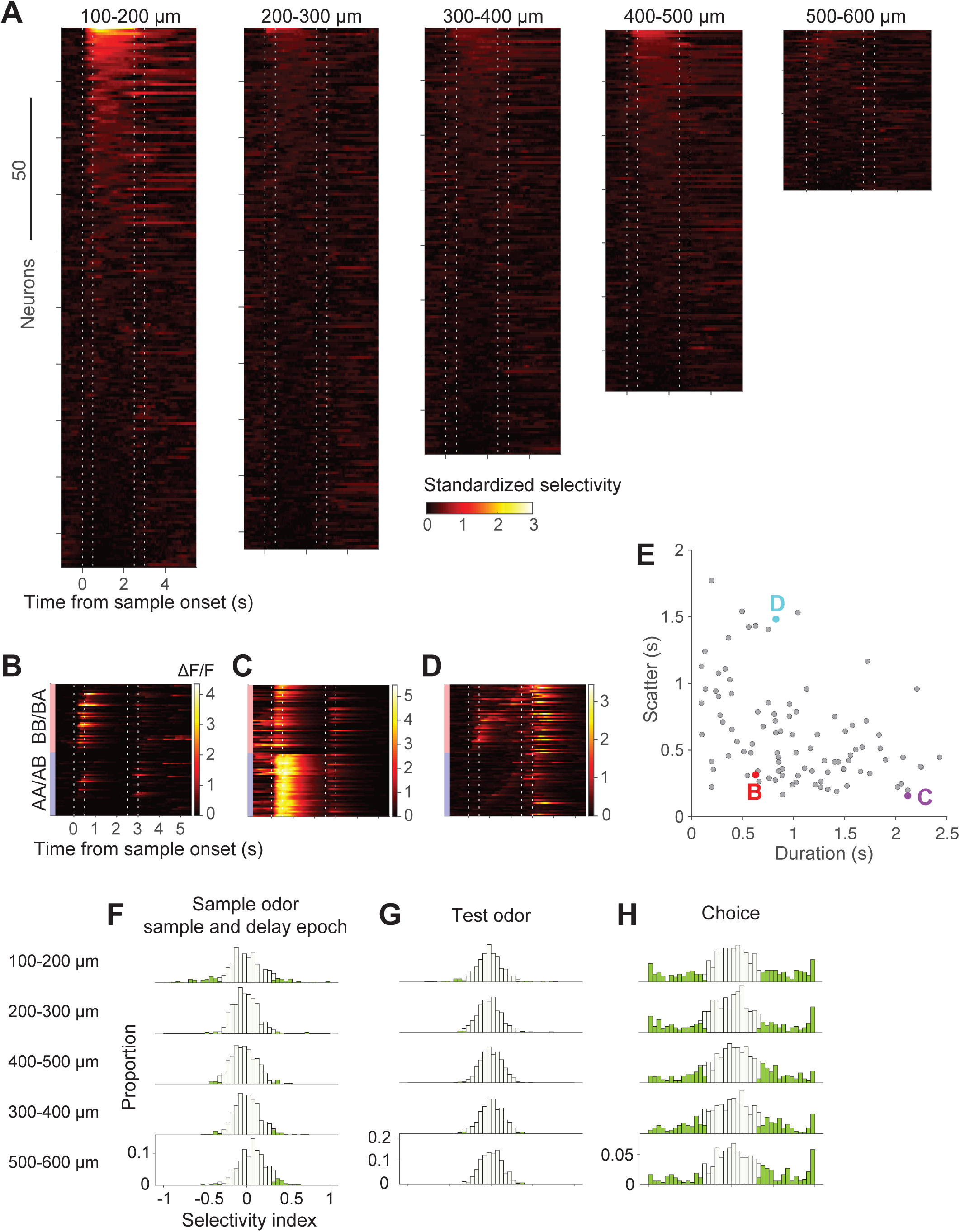
Two-photon calcium imaging of ALM. (**A**) Standardized sample odor selectivity of ALM neurons across five different cortical depths in an example animal. The standardized odor selectivity is the difference between the ΔF/F to odor A and B, normalized by the combined standard deviation (Methods). Each row is a neuron, sorted by the mean standardized selectivity during the sample and delay epochs. (**B-D**) ΔF/F traces of example sample-selective neurons. Each row is a trial, grouped by sample odor identity and sorted by peak response time to sample odor. (**E**) Time course of sample-selective responses. Each point shows the duration of the calcium transient (abscissa) against the scatter of the peak response (ordinate). Response duration is the median duration of each neuron’s response. Scatter is defined as the interquartile range of the peak response times across trials (Methods). The three colored points correspond to the neurons shown in panels (B-D). (**F-H**) Selectivity indices of neurons across five cortical depths for sample odor, test odor, and choice, respectively. n = 614, 514, 608, 512 and 379 neurons for the five cortical depths, respectively, from superficial to deep layers. Shading denotes statistical significance (*P* < 0.01, Mann-Whitney U test, two-tailed). See also Figure S5 and S6.

However, we also observed a large subset of neurons with striking odor selectivity during the sample and delay epoch (Fig. 5A, S5B, C). Interestingly, the odor selective calcium activity was heterogeneous in its timing during the sample and delay epochs, showing a variety of latencies and time scales (Fig. 5B-E). Some of the signals spanned the sample and part of the delay period, peaking at consistent times across trials (long duration, low scatter), spanning the sample and delay period (e.g., Fig. 5C). Other neurons exhibited more transient responses that occurred at different times across trials (short duration, high scatter; e.g., Fig. 5D). Although they are ordered on the graph for visualization, these brief, scattered activations appeared random. For example, their timing is uncorrelated in simultaneously recorded neurons (Fig. S6A, B). These characterizations also hold for the estimated spike rates, achieved through deconvolution of the raw Ca signals (Fig. S6C-F), despite a revision of the estimates of signal duration. From the imaging data, we adduce that a subpopulation of ALM neurons represents the identity of the sample odor through the sample and delay period.

These sample selective neurons appear to constitute a distinct cell type. They were concentrated in the superficial layer 2 (L2; 100-200 µm from the pial surface), where 18% responded selectively to one or the other the sample odor (Fig. 5A, F). They were only rarely encountered between 200 µm and 600 µm below the pial surface (*P* < 0.05, Kruskal-Wallis test with Tukey-Kramer multiple comparison). This layer specificity is not explained by an inability to measure calcium signals in deeper layers, as comparable fractions of choice-selective cells were found at all depths examined (*P* = 0.49, Kruskal-Wallis test; Fig. 5H, S5D). Indeed, a decoder, trained to classify the sample odor, performs much better using L2 neurons than using neurons from the other layers we sampled (Fig. 6A). In contrast, the capacity to decode choice is similar across all layers (Fig. 6B). Most of these sample selective neurons did not respond to the test odor (87 of 108, 81%; Fig. 6C) and only a few were choice selective or active during licking (Fig. 6D). Their time course distinguishes them from the rare odor selective responses encountered in our electrical recordings, and the absence of lick and choice activity distinguishes them from the dominant cell type in ALM.

**Figure 6:**
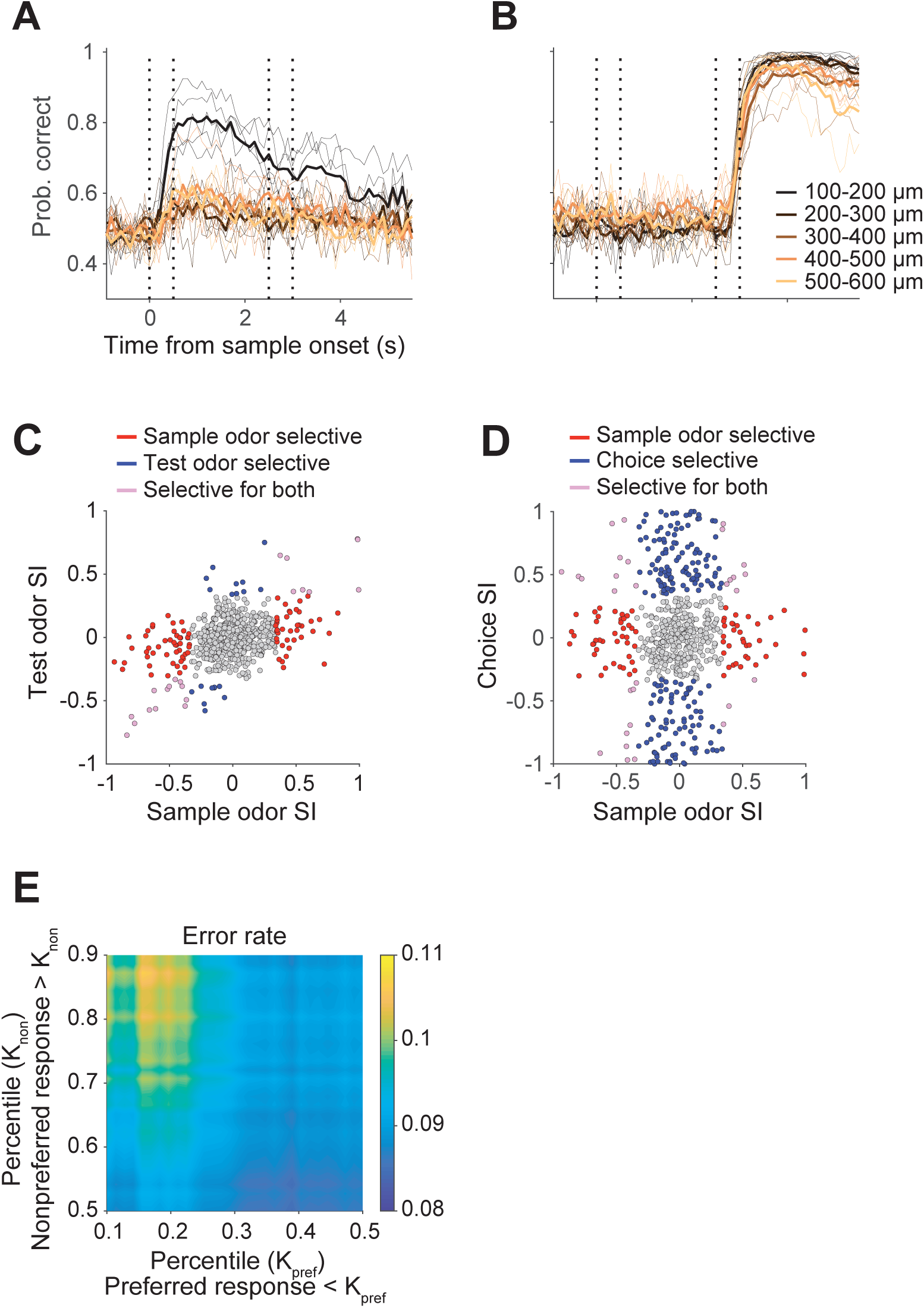
Sample-selective cells in ALM L2. (**A, B**) Performance of a support vector machine trained to classify the sample odor identity (A) or choice (B) using a subset of simultaneously imaged neurons from each of the five cortical depths. Each thin line represents the probability of correct classifications of the “held out” trials using data from one session (see Methods). Thick lines represent the mean of all sessions from a depth. (**C**) Selectivity indices of ALM L2 neurons for sample and test odors. Red, blue or purple denotes neurons selective for the sample odor, test odor or both, respectively. Non-selective neurons are shown in gray. (**D**) Selectivity indices of ALM L2 neurons for sample odor and choice. Red, blue or purple denotes neurons selective for the sample odor, choice or both, respectively. Non-selective neurons are shown in gray. (**E**) Trial-by-trial association between response and choice accuracy. The heatmap shows the fraction of errors when L2 neurons responded weakly to the preferred odor (*pref*) or strongly to the nonpreferred odor (*non*). The criteria for weak and strong are varied parametrically up to the median for *pref* (abscissa) and above the median for *non* (ordinate). The graph shows an increased probability of an error when the *pref* response is in the lower 20^th^ percentile or *nonpref* response is in the upper 30^th^ percentile. Logistic regression demonstrates that the effect is reliable across the dataset (*P* < 0.001; Methods).

The L2 neurons might resolve the perplexing inactivation result. ALM contains a representation of the sample odor throughout the delay period. It is therefore possible that the match/non-match decision is made within ALM, based on two external inputs that convey (*i*) the identity of the sample odor during the sample or delay period and (*ii*) and the identity of the test odor after the delay. Indeed, the activity of the L2 neurons during the sample and delay is informative about whether the mouse will ultimately succeed in the trial by making the correct choice. The logistic analysis (Fig. 6E) shows that when neurons selective for sample odor A, say, responded less to sample odor A or more strongly to sample odor B, the mouse was more likely to make an erroneous response. Such single trial correlations between neural response and choice, termed *choice probability*, have been exploited in perceptual decisions to support the proposal that a neuron’s response contributes to the decision process, either directly or via correlation with other neurons with similar response selectivity. In the present case the contribution is not to the match/non-match or lick left/right decision. Rather, it is a decision about which mapping to apply from test odors to lick responses (Fig. 1B).

### Potential mechanisms

We do not yet know the mechanism by which a representation of the sample odor in ALM affects the match/non-match decision, but it appears that some process requiring ALM in the sample and delay period must establish a state at the time of test that can implement the correct mapping between externally derived signals about the identity of the test odor and the activation of the appropriate lick neurons. Several possible mechanisms could underlie the formation of such a state. One possibility is that the state is defined by the persistent firing of sample-selective neurons as in standard “attractor” models of working memory (Fig. 7A) (Amit and Brunel, 1997; Goldman-Rakic, 1995). In this model, the requirement of ALM during the sample and delay epochs (Fig. 4) demands that ALM itself maintain a persistent representation of the sample odor. If the persistent sample representation is maintained in upstream areas but not ALM, the network model predicts that inactivating ALM would cause little behavioral impairment (Fig. 7B, top trace). This is inconsistent with our experimental observation. If the persistent sample representation is only maintained in ALM, the network model predicts that inactivation during the sample and early delay epochs should produce gross impairment (Fig. 7B, bottom trace). This too is inconsistent with the data (Fig. 4E). The weaker effects of inactivation when it was restricted to portions of the sample and delay epochs suggest that ALM’s representation of the sample odor can be partially recovered by activity in upstream areas. (Liu et al., 2014)(Liu et al., 2014)Thus, our experimental results argue for models in which multiple brain regions including both ALM and additional upstream areas each maintain a persistent representation of the sample odor (Fig. 7B, middle trace).

**Figure 7:**
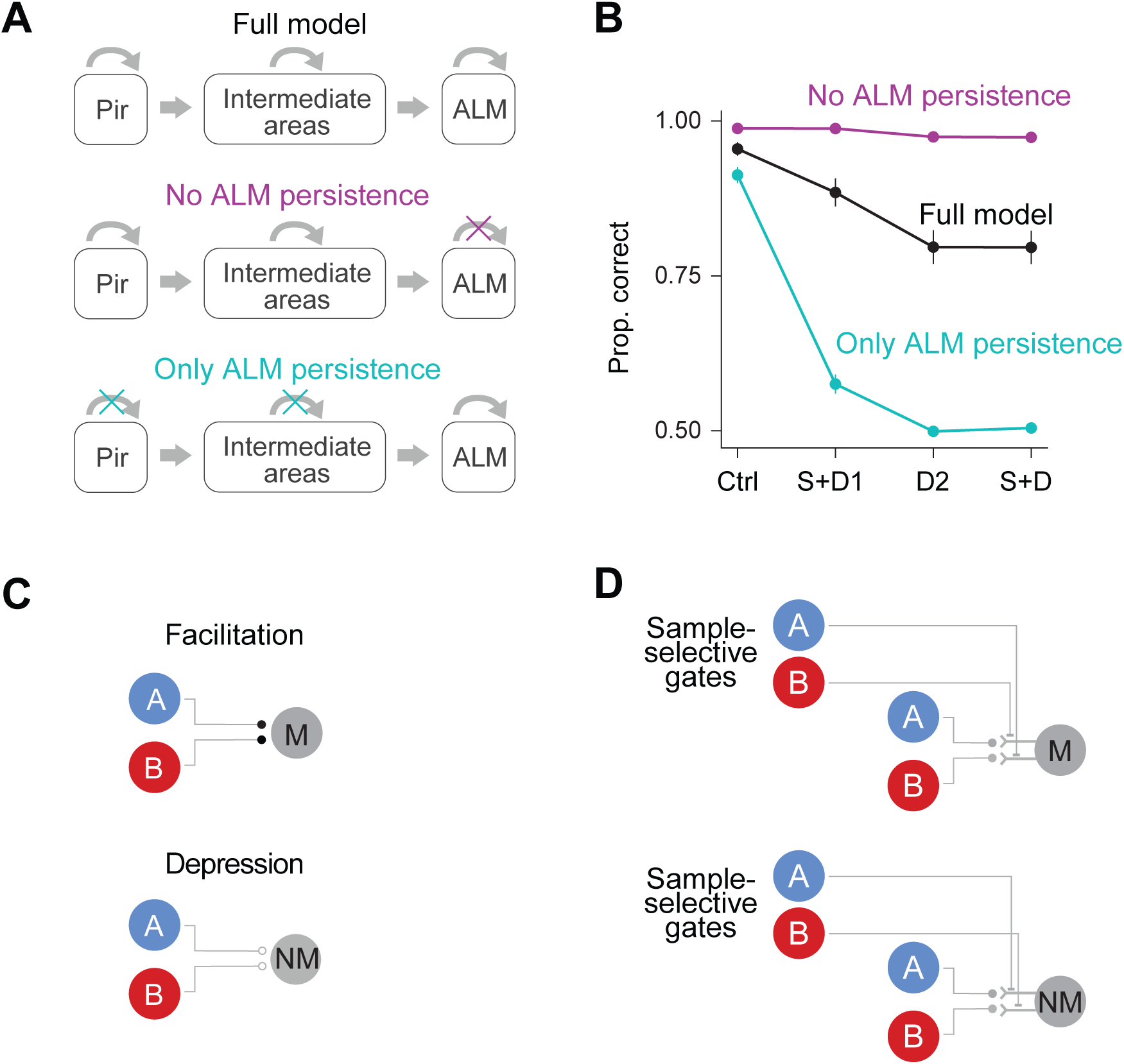
Neural models of the DMS task. (**A**) Three variants of the recurrent neural network model. Each contains three areas connected by fixed, random, feed forward projections (horizontal straight arrows). Top: all recurrent connections within each layer are trainable. Middle: recurrent connections within Pir and OFC, but not ALM, are trainable. Bottom: only recurrent connections within ALM are trainable. Note that recurrence within an area is necessary to achieve persistent activity that is not inherited from an upstream area. (**B**) Simulated effects of inactivating ALM during portions of the sample and delay epochs. Same conventions for the behavioral epochs as in Fig. 4E. (**C**) Synaptic facilitation and depression model. Facilitation leads to enhanced response to repeated presentation of the same stimulus in match selective neurons, whereas depression results in suppression in non-match selective neurons. We assume that the soma is inhibited during the sample epoch so that action potentials only occur after the test odor. (**D**) Dendritic gating model. Top: circuit supporting *match* (M). The sample odor suppresses excitatory inputs that convey the identity of the opposite test odor. Bottom: circuit supporting *nonmatch* (NM). A similar result could be achieved with presynaptic gating. This circuit also requires a mechanism to suppress spiking before the test epoch and a mechanism to reset the gate.

The attractor models posit that a representation of the sample odor is maintained through the persistent firing of neuronal assemblies. However, this memory might be maintained in other ways. We observed that impairment was more profound when ALM was silenced during the entire sample and delay epochs compared with inhibition only late in the delay (Fig. 4E). This suggests that a trace of the sample odor identity can persist in ALM during silencing. This trace may be maintained by facilitation or depression of synapses formed by axons of odor-selective neurons onto lick-left or lick-right neurons (Mongillo et al., 2008) (Fig. 7C). It could also be achieved through dendritic gating mechanisms that selectively route test odor information depending on the sample odor identity (Yang et al., 2016) (Fig. 7D). In this model, neurons selective for a particular sample odor enhance or suppress dendritic branches of lick-selective neurons where either A or B inputs concentrate. This synaptic modulation could be controlled by sample-selective neurons in ALM layer 2.

## Discussion

Decisions that engage cognitive capacities entail at least three elemental processes: (i) the maintenance of information over time scales that can extend for seconds prior to a response; (ii) the association of the same information with different responses; and (iii) the association of the same response with different information. The first process requires planning and working memory (Fuster, 1973; Goldman-Rakic, 1995; Romo et al., 1999), whereas the latter two involve flexible, context dependent routing of information to appropriate outputs.

Our findings suggest an unexpected role of ALM in processing perceptual information. Neurons in ALM are known to play an essential role in the planning of licking (Svoboda and Li, 2018). In primates and more recently in rodents, preparatory activity in the premotor parietal circuit has been shown to play a role in perceptual decisions in which attributes of sensory stimuli (e.g., the direction of moving random dots) instruct the selection of a movement (Hanks et al., 2015; Shadlen and Newsome, 2001). In this behavioral paradigm, movement selection evolves gradually and reflects the deliberation process leading to a decision (Gold and Shadlen, 2003; Selen et al., 2012; Spivey et al., 2005). These observations might suggest that premotor area would play a role in the perceptual decision in our DMS task. However, in our task a plan to lick left or right cannot be prepared until the test odor arrives. What can be prepared is the sensory-motor association between the test odor and the appropriate lick direction. Therefore, the impairment in performance arising from the inactivation of ALM does not reflect a disruption of movement preparation but of the state of the circuit that readies it to process information about the test odor and select the appropriate lick response.

Previous studies have demonstrated persistent activity in ALM in association with perceptual decision making. For example, elevated activity was reported in direction selective lick neurons when mice associated different haptic stimuli (applied to a whisker) with a lick to the left or right (Chen et al., 2017; Guo et al., 2014; Li et al., 2015; Svoboda and Li, 2018). This persistent activity is involved in planning and driving movement based on a sensory-response association. In our study, on the other hand, persistent activity in sample-selective neurons represents a sensory category rather than a plan of action, because the action cannot be selected until after the delay period upon receipt of the test odor. Hence, unlike in a sensory-instructed delayed response task (Guo et al., 2014; Li et al., 2015) in which persistent activity represents decision outcome, the activity of our L2 neurons is not the outcome of the DMS decision because this decision has yet to be made. This activity could be viewed as representing an intermediate decision about which mapping from test odor to lick response to employ (Fig. 1B). It is possible that our sample selective neurons overlap those reported previously, but we suspect the overlap is minimal. The persistent activity we observed was concentrated primarily in superficial L2 neurons that did not exhibit lick responses, whereas the persistent activity reported in the somatosensory instructed delay task was found in all cortical layers and in many neurons that responded to error responses in a way that agrees with their role in motor planning.

We considered several alternative explanations of the impairment produced by ALM inactivation. The control experiments rule out a simple motor impairment or motor bias, allowing us to focus on the delay period activity. Our interpretation is that the representation of the sample odor in ALM establishes a state of the ALM circuit such that it can perform a simple sensory-response association at the time of test. An alternative is that ALM must prepare both potential motor plans during the delay period (Cisek and Kalaska, 2005) in order for one to be selected by an instruction from another brain area, and it is solely the disruption of this preparatory activity that explains the impairment. This account predicts incorrectly that inactivation would also impair performance on the A × A/B and C × C/D tasks. It is also inconsistent with the neural recordings, which failed to reveal simultaneous preparation of the two lick directions during the delay period.

Another alternative explanation is that inactivation before the test period disrupts a general state of readiness in ALM to respond to any signal instructing an action to be performed. We find this unlikely, because to account for our behavioral observations, this state would need to be specific to the DMS task and not control tasks with identical temporal structures and actions. Finally, we note that DMS is a more difficult task than the controls. Therefore, it may be argued that DMS is more susceptible to perturbations. However, it is not the motor action that makes DMS more difficult than the controls, because all the tasks require the identical motor responses. If the only role of ALM is motor preparation, then inactivating ALM should not result in differential impairment of the DMS and control tasks (Fig. 4H). More importantly, our perturbation affected ALM before a decision could be prepared, because the decision cannot be made before the test odor arrives. Previous studies of decisions to lick to the left or right examined only the preparatory phase after the decision was instructed (Guo et al., 2014; Li et al., 2015).

Neurons in superficial layers of the neocortex receive input from controlling structures such as intralaminar and accessory thalamic nuclei (Jones, 1998) as well as long range feedback, e.g., perirhinal cortex to ALM (Zingg et al., 2014). These layers contain the distal dendrites of projection neurons in deeper layers, and activation/suppression of these dendrites can alter the functional properties of the projection neurons (Bittner et al., 2017; Larkum et al., 1999, 2009; Polsky et al., 2004). Thus, it is possible that the selectivity to the sample odor that we observed in L2 neurons might reflect a general mechanism that alters the state of ALM circuitry to flexibly implement different sensorimotor mappings. It remains to be seen if the L2 neurons are in fact critical and if so, how they alter the state of the ALM circuit. However, the association between the probability of an error and the level of activity on single trials (Fig. 6K) suggests they play a functional role.

It may seem counterintuitive that a premotor area would play a critical role in a decision about the relationship between two sensory stimuli. However, the ultimate goal of both sensory and motor systems is not to identify stimulus features or categories, but to determine whether to act one way or another. A potential action might require support from different sources of sensory input. Thus, decisions to act—even if only provisionally—rest on a capacity of the motor system to establish functional connections that allow it to integrate different sources of sensory information, and this information infiltrates the responses of neurons in cortical areas that are appropriately designated premotor or associative (Cisek, 2011; Shadlen and Kiani, 2013). These functional connections must be rapidly modifiable as the demands of an organism and its context change. This capacity for dynamic circuit configuration is likely to support a wide range of cognitive phenomena involving flexible routing, decision making, and perceptual inference (Shadlen and Shohamy, 2016).

## Acknowledgements

We thank T. Harris for the acute H3 probes; T. Tabachnik, D. Peterka, G. Johnson, J. Wang, J. Goldbas, A. Baker and M. Chin for technic assistance. K. Svoboda, Z. Guo, L. Abbott, A. Losonczy, R. Bruno, C. Rodgers, C. Lacefield, K. Shin, and B. Hu for advice on experiments and analysis; D. Wolpert and members of the Axel and Shadlen labs for helpful discussions; C. Eccard for assistance in the preparation of the manuscript; M. Myles for figure artwork. Z.W. is a Simons Junior Fellow. A.L.-K. is supported by NSF NeuroNex Award DBI-1707398, the Burroughs Wellcome Foundation, and the Gatsby Charitable Foundation. R.A. and M.N.S. are supported by the Howard Hughes Medical Institute (HHMI). R.A. is also supported by the Simons Foundation.

## Author Contributions

Z.W., R.A. and M.N.S. conceived and designed the study. Z.W. performed all the experiments with assistance from P.S. and A.T. A.L.-K. implemented the attractor network models. Z.W., A.L.-K., R.A. and M.N.S. interpreted the results and wrote the paper.

## Data and materials availability

The data, code, and materials that support the findings of this study are available from the corresponding authors upon reasonable request.

---

**Figure S1:**
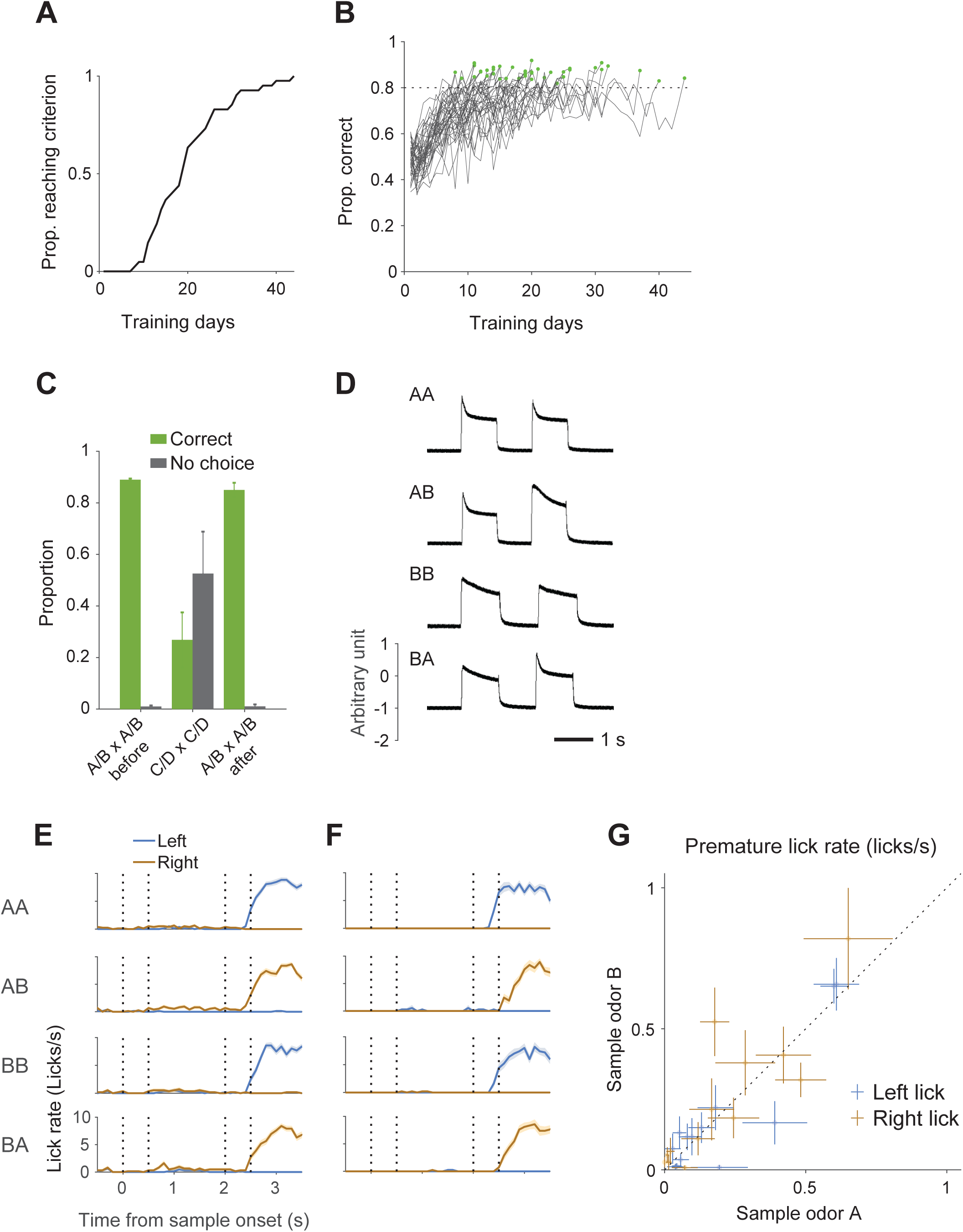
Mouse training on the DMS task. Related to Figure 1. (**A**) Cumulative proportion of animals that reach training criterion as a function of training days (n = 41). Training days only include the days in which mice were presented with all four trial types. (**B**) Progression of each mouse’s performance on the DMS task. Each line represents the data from one animal; the green dot indicates the day on which they reached criterion performance. (**C**) Performance of animals on a set of novel odors, C/D × C/D. The animals (n = 5) were previously trained to criterion on the A/B × A/B task. The poor performance is characterized by a smaller proportion of correct choices and a large proportion of “no choice” trials. (**D**) Time course of the sample and test odor delivery as measured by a photo-ionization detector. The measurements were taken in the training phase when sample and test epochs were 1 s. (**E**, **F**) Premature licks are indistinguishable for the two sample odors. Lick rates of two example animals, grouped by trial type. Only correct trials are shown. The lines and shadings indicate mean and s.e.m. (n = 47-79 trials). (**G**) Mean lick rates during the sample and delay epochs for trials in which the sample odor was A, plotted against the rates for sample odor B. Each point represents data from one animal (n = 13). Error bars are s.e.m.

**Figure S2:**
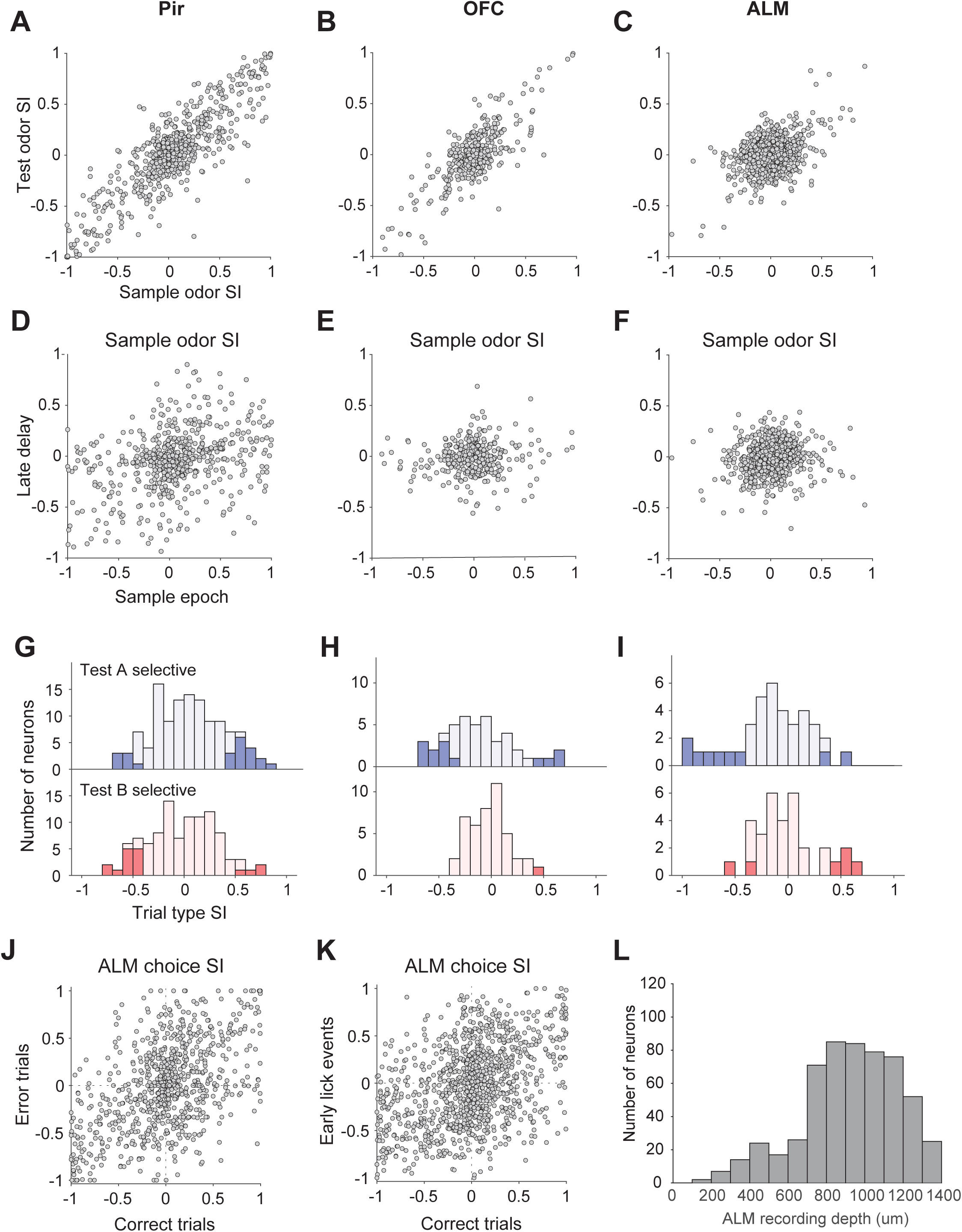
Selectivity of task-related variables in Pir, OFC and ALM and the depth of ALM neurons. Related to Figure 2. The three brain areas correspond to the left, middle and right columns of panels (A-I). (**A-C**) Selectivity indices of Pir, OFC, and ALM neurons for sample and test odors (see Methods). Pearson’s correlation *R* values: Pir, 0.86 (*P* = 1.6e-184); OFC, 0.79 (*P* = 1.3e-82); ALM, 0.41 (*P* = 5.9e-45). (**D-F)** Selectivity indices of neurons for sample odor in the sample epoch and in the late delay. Pearson’s correlation *R* values: Pir, 0.34 (*P* = 9.5e-19); OFC, 0.076 (*P* = 0.14); ALM, 0.12 (*P* = 5.7e-5). (**G-I**) Selectivity indices of neurons for trial type (see Methods). Only test odor selective neurons are included. Top: selectivity indices of test odor A-preferring neurons for sample odor identity; a positive value indicates a preference for AA trials and, a negative value indicates a preference for BA trials. Shading denotes statistical significance (Mann-Whitney U test, *P* < 0.01, two-tailed). Bottom: same for test odor B-preferring neurons. (**J**) Choice selectivity indices of ALM neurons in correct and error trials. Pearson’s correlation, *R* = 0.51, *P* = 4.6e-53. (**K**) Choice selectivity indices of ALM neurons in correct trials and in early lick events. *R* = 0.42, *P* = 4.6e-47. (**L**) The depth of the recorded ALM units measured from the pial surface.

**Figure S3:**
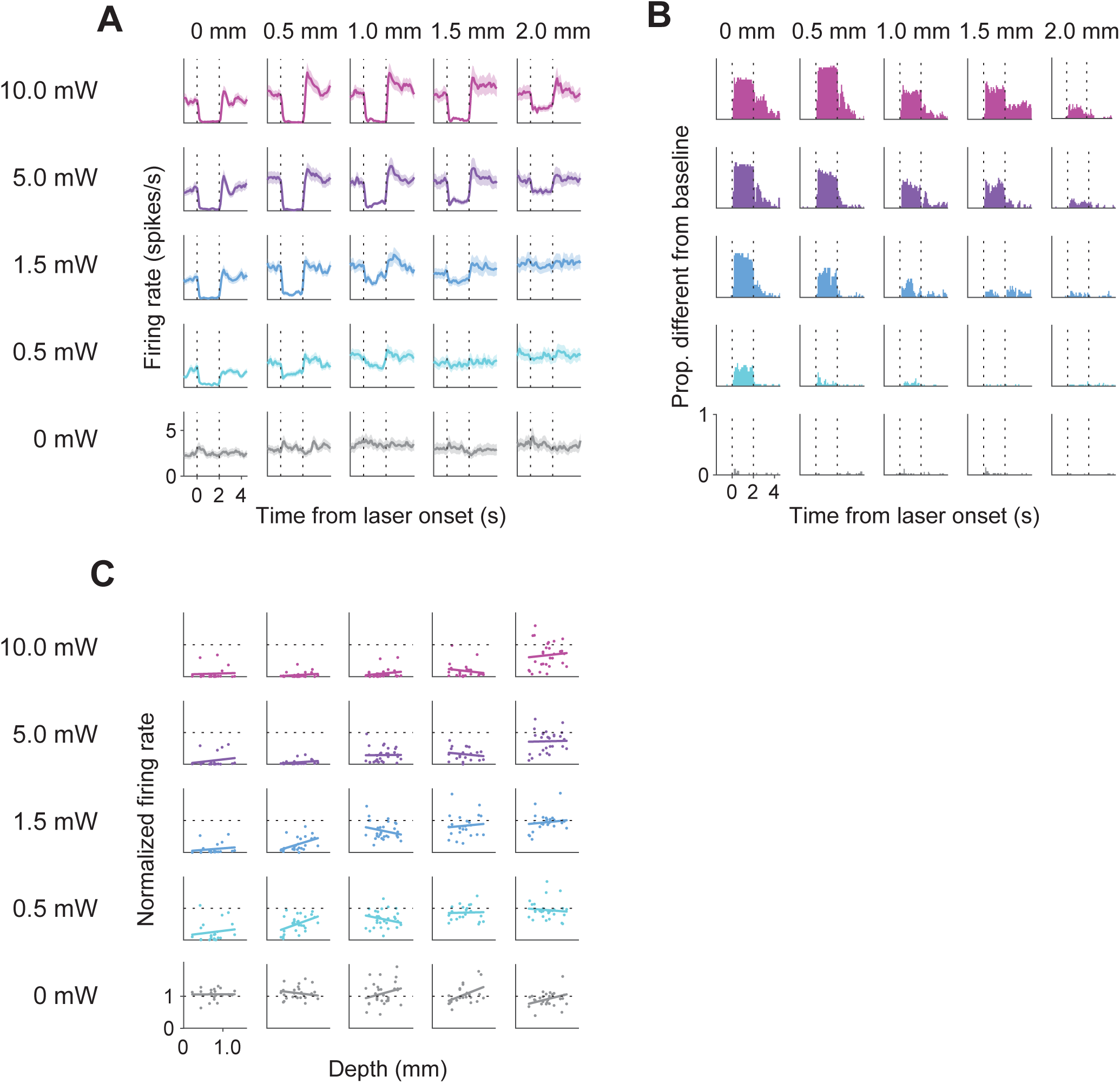
Calibration of laser power for photoinhibition. Related to Figure 4. (**A**) Mean firing rate of ALM neurons over a 2 s period of photoinhibition (n = 74). Shading indicates s.e.m. The rows denote different laser powers and the columns indicate the distance of the recording sites from the center of the laser spot along the cortical surface. The 2 s illumination includes a power ramp-down in the final 250 ms. (**B**) Proportion of neurons with firing rates that are significantly different from baseline (Mann-Whitney U test, *P* < 0.01). (**C**) Normalized mean firing rate of neurons during photoinhibition, plotted as a function of cortical depth from the pial surface. The column labels are the same as in (A). Each point represents data from one neuron. The lines are linear fits. The firing rates were normalized to the average response in a 1s baseline period before laser onset. Only neurons with baseline firing rate > 1 spikes/s were included.

**Figure S4:**
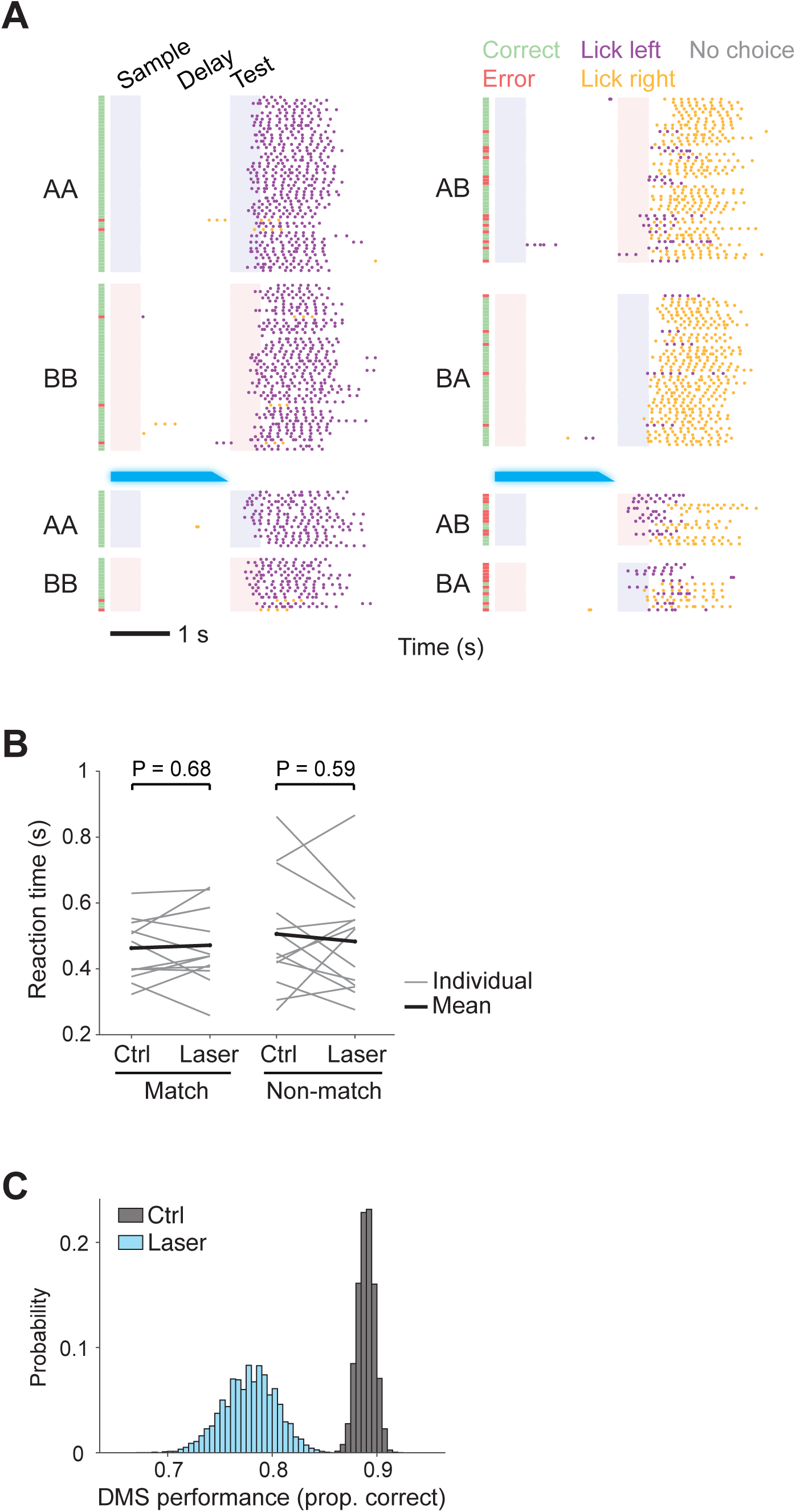
Licking and side bias in ALM inactivation. Related to Figure 4. (**A**) An example behavioral session with and without ALM inactivation. The animal still licked to the two ports but made more errors. The top two rows are control trials and the bottom two rows are inactivation trials. All the trials were randomly presented but grouped here for plotting. The blue bars indicate the timing of the photoinhibition. (**B**) Reaction time in match and non-match trials with and without ALM inactivation. Each thin line represents data from one animal; the thick lines connect the means from 13 animals. Control and laser groups were compared with paired *t*-test. (**C**) Estimation of behavioral impairment not accounted for by a side bias. Monte Carlo methods were used to estimate the change in accuracy that was not attributable to a bias. We simulated datasets matched in size to the experiments in Fig. 4B, using the logistic fits to the data but with the laser induced bias set to zero (*β_3_* = 0, Eqs. 7 and 8, see Methods). The histograms show distributions of Proportion Correct for control and laser from 10,000 simulations. The difference in expectations, 0.89 ± 0.008 (mean ± standard deviation) to 0.78 ± 0.025 correct (*P* = 1.8e-5, *t*-test), is a conservative estimate of the impairment that is not induced by a laser induced bias. The estimate is conservative because bias could change as a consequence of uncertainty associated with impairment on the task. The simulation complements the logistic regression analysis in which choices are coded to maximize the effect of laser induced inactivation on bias (Eqs. 7 and 8). Estimated coefficients are *β_1_* = 2.10 ± 0.059, *P* = 4.0e-281; *β_2_* = 1.00 ± 0.11, *P* = 7.3e-19; *β_3_* = −0.84 ± 0.11, *P* = 1.5e-13. Note that sensitivity on the task is significantly impaired by inactivation (*β_3_ <* 0).

**Figure S5:**
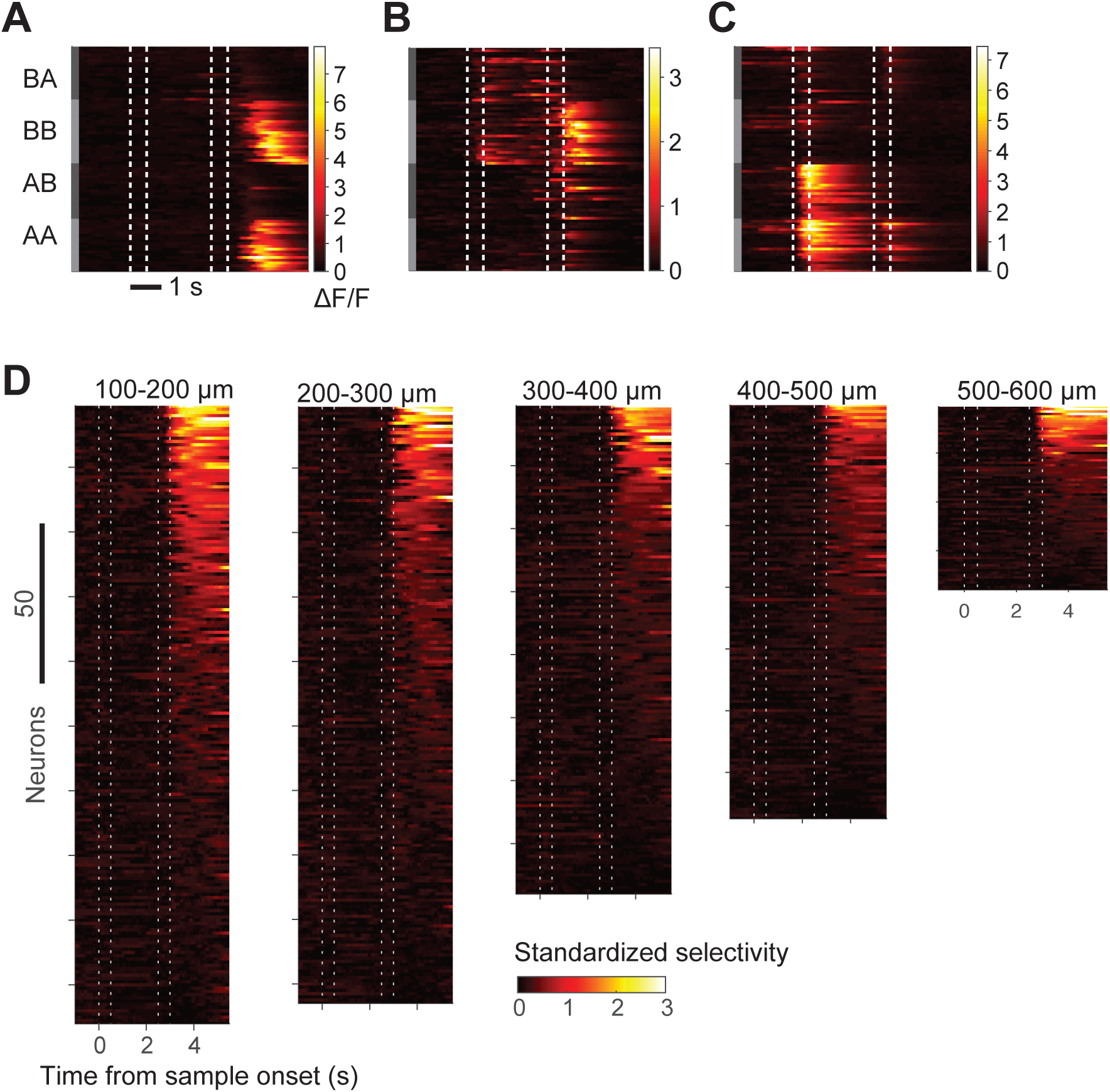
Calcium responses and selectivity of ALM neurons. Related to Figure 5. (**A-C**) ΔF/F of three example ALM neurons. Each row is a trial, grouped by trial type. (**D**) Standardized choice selectivity of ALM neurons across different cortical depths (see Methods). Each row is a neuron, sorted by the mean standardized choice selectivity within 2.5 s of test onset.

**Figure S6:**
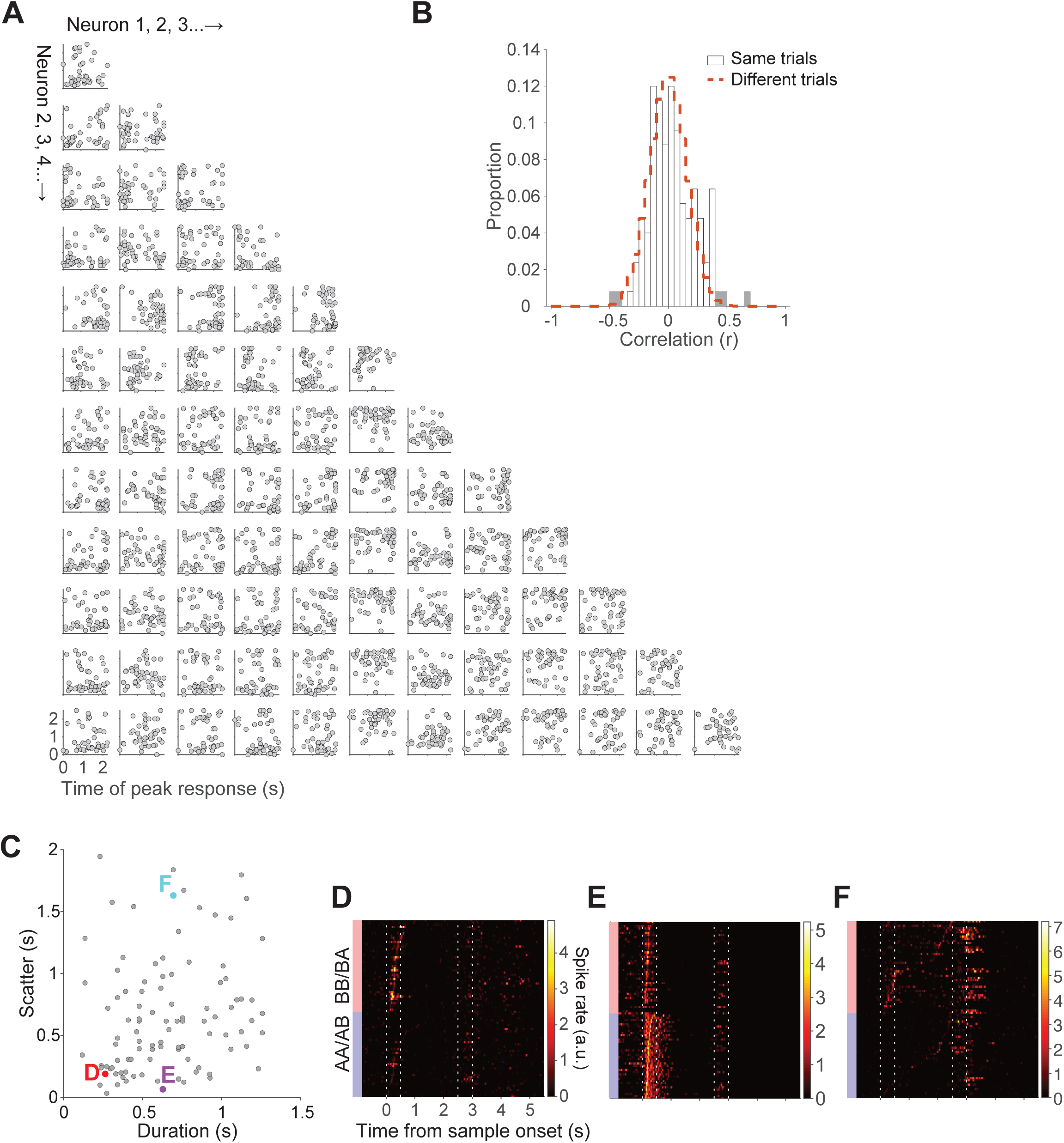
Correlation of peak response time and deconvolved spike rates. Related to Figure 5. (**A**) Correlation of the peak response time between simultaneously recorded neuron pairs from a session. Only sample odor B selective neurons with a peak time scatter (see Fig. 6E) over 0.6 s are included. (**B**) Distribution of the correlation coefficients of the peak response from all neuron pairs. The shading indicates coefficients that are significantly different from chance (*P* < 0.01; Fisher z, uncorrected for multiple comparisons). The red dashed curve shows the distribution of correlation coefficients under the null hypothesis (estimated from a shuffle control; see Methods). (**C**) The duration and scatter of L2 sample-selective neurons using deconvolved spike rates. The three colored points correspond to the neurons shown in panels (D-F). (**D-F**) The deconvolved spike rates of three example neurons shown in Fig. 6B-D, respectively. Each row is a trial, grouped by sample odor identity and sorted by peak response time to sample odor.

## STAR Methods

### Key Resources Table

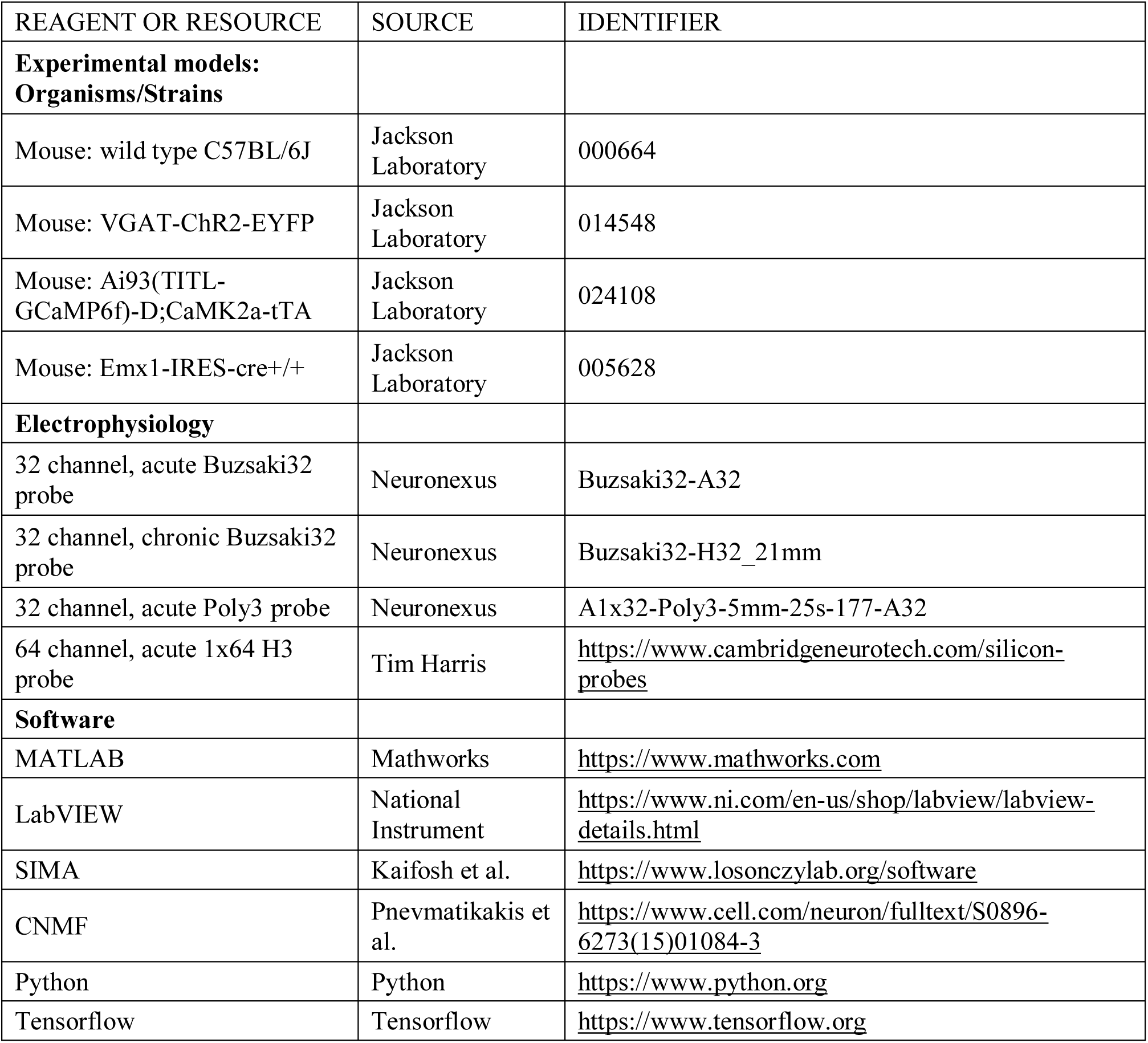

### Animals

Animals were maintained on 12h: 12h light/dark cycle with food and water available *ad libitum*. Animal care and experiments were carried out in accordance with the NIH guidelines and approved by the Columbia University Institutional Animal Care and Use Committee (IACUC).

Mice were water-restricted during the training and testing phases. Experimental sessions were 1-2 hours, during which mice received 0.5-1.5 ml of water. Animals received supplemental water as necessary to maintain their body weights. Aseptic surgeries were carried out under ketamine (100 mg/kg)/xylazine (10 mg/kg) or 1-3% isoflurane anesthesia. Buprenorphine (0.05-0.1 mg/kg) and carprofen (5 mg/kg) were administered for postoperative analgesia.

This study is based on data from 41 mice (both males and females, 2-8 months old). 5 C57BL/6J and 6 VGAT-ChR2-EYFP (Jackson Laboratory, JAX 014548) mice were used for electrophysiology recording. 2 untrained VGAT-ChR2-EYFP mice were used to characterize the inhibition at different laser powers. 29 VGAT-ChR2-EYFP mice were used for inhibition experiments, 6 of which were also used for simultaneous recording during inhibition. Two-photon imaging data were collected from 5 Emx1-cre+/-; TITL-GCaMP6f+/-; CaMK2a-tTA+ mice. These mice were created by crossing Ai93(TITL-GCaMP6f)-D;CaMK2a-tTA (JAX 024108) to Emx1-IRES-cre+/+ (JAX 005628) line.

### Behavior training

Before training, mice were implanted with a custom-made titanium head post (Guo et al., 2014b). The scalp and periosteum over the dorsal surface of the skull were removed, and a head post was placed on the skull, aligned with the lambda suture and cemented in place with C&B metabond (Parkwell, S380). After at least 2-3 days of recovery, animals were water-restricted and accustomed to head-fixation following procedures described in Guo et al., 2014 (Guo et al., 2014b) and then trained on a custom-assembled apparatus.

Odorants were delivered with a custom-made olfactometer and custom-written LabVIEW programs (National Instruments). (+)-α-Pinene (odor A, Sigma-Aldrich, 268070), cis-3-Hexen-1-ol (odor B, Sigma-Aldrich, W256307), (R)-(+)-Limonene (odor C, Sigma-Aldrich, 183164), and Methyl butyrate (odor D, Sigma-Aldrich, 246093) were chosen for their lack of innate valence to mice (Root et al., 2014) and their low adhesion to the surface of the olfactometer. The odorants were diluted 100-fold in mineral oil (Fisher Scientific, O121-1) and then loaded on syringe filters (GE healthcare, 6888-2527 or 6823-1327). The air flow was maintained at 1.0 L/min. We confirmed the rapid kinetics (Fig. S1D) of these odorants with photo-ionization detector (Aurora Scientific, 200B). Over the course of a behavioral session, the odorants on the syringe filters gradually deplete. Thus animals were unlikely to rely on the absolute concentration of the odorants which changed constantly, but rather the identity of the odors, as evidenced by the stable sensory responses in Pir over a session (Fig. 2A).

During training, mice were presented with a sample odor (1.0 s duration) and a test odor (1.0 s) separated by a delay epoch (1.0 s). After hearing an auditory “go” cue (0.1 s, 5 KHz pure tone), they were free to report their decision by licking to one of the two syringe ports positioned in front of their mouth, and collect a water reward at the same port if they were correct. Many animals started licking in the test epoch, which was permitted, but only the licks during the 2 s “response window” following the “go” cue were considered their choice. To prevent mice from “probing” for the correct port by rapidly switching between the two ports, we required mice to commit to their choices. A choice is scored as correct only when the first two licks were on the correct port. If they first licked on the incorrect port, that trial was scored as an error. If they licked once on the correct port and then on the incorrect port, that trial was also scored as an “error”. If they did not lick during the “response window” or only licked once on the correct port, that trial was scored as “no choice”.

The two odorants give rise to four unique pairs of sample and test odors (AA, AB, BB, and BA), or trial types, and were randomly presented in each session. The match trials (AA, BB) were rewarded on the left port and non-match trials (AB, BA) on the right. Mice were punished by a brief timeout (3-8 s) when they made an error. “No choice” trials were rare and typically occurred in the very early training stage or at the end of a session when the mice were sated. Animals completed this training stage when they achieved a criterion of at least 80% correct for each trial type in a single session. They then underwent additional training to suppress “premature” licking before the test epoch. Such early licks were punished by an immediate 0.1-0.2 s siren (RadioShack, 273-079), followed by a 0.5-1.0 s pause in that trial, and a longer inter-trial interval. We required the proportion of trials with premature licking to be less than ∼20%. After achieving these milestones (median 25 days; IQR 21-31 days) the sample and test durations were reduced from 1.0 s to 0.5 s, and the delay was increased from 1.0 s to 1.5-4.0 s depending on the experiment (1.5 s in extracellular recording, 2.0 s in imaging experiments, 1.5, 2.5 or 4.0 s in optogenetic inactivation). Mice performed at 90% correct (interquartile range 88-92%) when they entered the testing phase.

### Electrophysiology

Extracellular recordings were made acutely or chronically in head-fixed animals using 32- or 64-channel silicon probes (Buzsaki32 and Poly3, NeuroNexus; 1×64 acute H3 probe, HHMI). The probes were targeted stereotaxically to Pir, OFC and ALM using Bregma coordinates as follows: Pir: AP 1.9-2.4 mm, ML 1.5-2.1 mm, DV 2.3-2.9 mm; OFC: AP 2.2-2.8 mm, ML 0.7-1.2 mm, DV 1.2-1.8 mm; ALM: AP 2.5 mm, ML 1.5 mm, DV 0-1.1 mm.

In acute recordings, a small craniotomy (0.3-0.8 mm in diameter) was made over the targeted area before the recording session. Recording depth from the pial surface was inferred from micromanipulator reading. After each recording session, the brain surface was covered with silicone gel (3-4680, Dow Corning) and Kwik-Sil (World Precision Instruments). In chronic recordings, the probe (Buzsaki32-H32_21mm, NeuroNexus) was attached to a custom-made microdrive which allows for advancement of the shanks. The microdrive was then implanted and cemented with C&B metabond and dental acrylic (Lang Dental Jet Repair Acrylic, 1223CLR). We advanced the shanks by 50 µm per day at the end of each recording session.

### Two-photon calcium imaging

Calcium imaging was performed on Emx1-cre+/-; TITL-GCaMP6f+/-; CaMK2a-tTA+ mice. A square craniotomy (2mm side) was made above left or right ALM, along the superior sagittal sinus and the inferior cerebral vein. The imaging window was constructed from three stacked layers of custom-cut coverglass (CS-3S, Warner Instruments) and cemented with C&B metabond. Animals were allowed 1-2 weeks of recovery before the imaging sessions began. Images were acquired with a Bruker Ultima two-photon microscope under resonant galvo scanning mode. The light source was a femtosecond pulsed laser (Chameleon Vision II, Coherent). The objective was a 16X water immersion lens (Nikon, 0.8 NA, 3mm working distance). GCaMP6f was excited at 920nm and images (512 x 512 pixels, ∼820 µm x 820 µm field of view) were acquired at ∼30 Hz.

### Photostimulation

Animals were prepared with a clear skullcap to achieve optical access to ALM (Guo et al., 2014a). Briefly, after removing the scalp and periosteum over the dorsal surface of the skull, a layer of cyanoacrylate adhesive (Krazy glue, Elmer’s Products Inc.) was directly applied to the intact skull. The entire skull was then covered with a thin layer of the clear dental acrylic (Lang Dental) with a head post cemented over the lambda suture. Before photostimulation sessions, the dental acrylic was polished (0321B, Shofu Dental Corporation) and covered with a thin layer of clear nail polish to reduce glare (Part No. 72180, Electron Microscopy Sciences). Light from a 473nm laser (MLL-FN-473-50mW, Ultralasers, Inc.) was directed to an optic fiber and split into two paths (FCMH2-FCL, Thorlabs, Ø200 µm Core, 0.39 NA). The two optic fibers were positioned over ALM on each hemisphere. The light transmission through the skullcap is ∼50% in average power, as measured directly with a light meter (PM100D, Thorlabs) with freshly dissected skullcap, consistent with previous measurements (Guo et al., 2014a).

We used 40 Hz photostimulation with a sinusoidal temporal profile (3 mW average power as light reaches the skull, ∼1.5 mW on the brain surface). The photoinhibition inactivated a cortical area of ∼1mm radius, as the population firing rate drops to ∼50% at 1mm away from the center of the laser beam (Fig. S3A). To reduce rebound excitation after laser offset, we included a 250 ms linear power ramp-down at the end of the photostimulation (Fig. 4A) unless otherwise indicated. In the interleaved A/B × A/B & C × C/D experiment, the delay epoch was extended to 4s while sample and test epochs were kept at 0.5 s each. Here we used a 500 ms ramp-down at the end of the 4s photostimulation, which terminates 500 ms before the test odor onset to allow more recovery time. To prevent the mice from distinguishing photostimulation trials from control trials, a masking flash (40 Hz sinusoidal profile) was delivered with 470 nm LED (Luxeon Star) and LED driver (SLA-1200-2, Mightex Systems) in front of the animals’ eyes on all trials. The masking flash was not phased locked to the photostimulation, began at sample onset and lasted until the end of test, covering the entire stimulus and delay epochs in which photostimulation could occur.

For the experiments in which we varied the duration of inactivation (Fig. 4E), photostimulation was either limited to (*i*) the 0.5s sample epoch and the first 1.5s of the 2.5s delay epoch, or (*ii*) the last 1.0s of the delay epoch, or (*iii*) the entire sample plus delay epoch. These inactivation trials were randomly interleaved to constitute 25% of all trials. For each animal, multiple (2-3) behavioral sessions were pooled to collect at least 10 trials for each of the four trial types and the three inactivation durations.

### Simultaneous photostimulation and recording

We calibrated the laser power for ALM and OFC inactivation by recording from these two areas during photoinhibition in awake animals. For ALM inactivation, we positioned an acute 1×64 H3 probe at various distances from the optic fiber over ALM (0-2.0 mm in 0.5 mm increments). A range of laser powers (0.5 mW, 1.5 mW, 5.0 mW, and 10.0 mW) as well as controls were examined at each location. We chose 1.5 mW power on the brain surface as it inactivates a cortical area of ∼1mm radius and produces minimal rebound activity after laser offset (Fig. S3). For OFC inactivation, an optic fiber was targeted to Bregma AP 2.5 mm, ML 1.0 mm, DV 1.4 mm. The recording probe was then positioned at AP 2.5 mm, ML 1.5 mm, DV 1.0-2.3 mm, where it is close to the border of OFC with agranular insular cortex. We recorded the neural responses similarly at a range of laser powers and chose 1.0 mW to silence OFC while minimizing the impact on neighboring brain areas.

For simultaneous recording and inactivation when animals performed the DMS task, we used the clear skullcap preparation for photoinhibition and made a small craniotomy lateral to the optic fiber to insert the probe. The probe was advanced at 60-70° angle from the horizontal plane at Bregma AP 2.3-2.5 mm, ML 2.0-2.5 mm to record from ALM, OFC, and Pir at different depth.

### Electrophysiology data analysis

The 32- or 64-channel recording data were digitized at 40 KHz and acquired with OmniPlex D system (Plexon Inc.) The voltage signals were high-pass filtered (200 Hz, Bessel) and sorted automatically with Kilosort (Pachitariu et al., 2016). The clusters were then manually curated with Phy GUI (Rossant et al., 2016) to merge spikes from the same units and to remove noises and units that were not well isolated. Recording depths were inferred from micromanipulator readings in acute recordings or microdrive turns in chronic recordings.

We determined sample odor selectivity for each neuron by comparing the spike counts during the sample epoch (0.1-0.6 s from sample onset) or late delay (1.5-2.0 s from sample onset) between sample odor A group (AA, AB) and sample odor B group (BA, BB). The 0.1 s time offset from the sample epoch accounts for the olfactometer valve time. A neuron is considered odor selective if its responses to sample odor A and B are significantly different by a two-tailed Mann-Whitney U test (*P* < 0.01, not corrected for multiple comparisons). The selectivity index (SI) was computed as follows for each unit: First, the trial-by-trial spike counts from the responses to odor A and B were used to construct a Receiver Operating Characteristic (ROC). The area under the ROC (AuROC) is the probability that a randomly sampled response associated with odor A is larger than a randomly sampled response associated with odor B. The selectivity index is

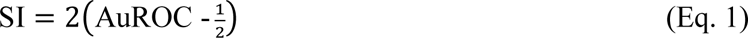

so that ±1 represents perfect discriminability (i.e., no overlap of the response counts) and 0 indicates chance-level separation. Positive SI connotes a preference for sample odor A. We determined selectivity for test odor and choice in a similar fashion by comparing spike counts during the test epoch (2.1-2.6 s from sample onset), with appropriate trial grouping. Positive SI for choice connotes a preference for licking to the left spout.

The trial type selectivity in Fig. S2G-I was computed as follows. Neurons from each of the three areas were first selected by their test odor selectivity. Only neurons that are test odor selective as determined by a Mann-Whitney U test (*P* < 0.01, not corrected for multiple comparisons) were included in the subsequent analysis. These test odor selective neurons were then treated separately based on their preference for test odor A or B. If a neuron prefers test odor A, its selectivity index for trial type was computed based on its responses to AA vs. BA trials, in a similar fashion described above. Positive SI connotes a preference for AA. If a neuron prefers test odor B, its trial type selectivity is computed based on its response to AB vs. BB, and positive SI connotes a preference for AB.

Graphs depicting population odor selectivity of a brain area show the difference in firing rates associated with the preferred and nonpreferred odors (Fig. 3A, 4I-N). The preferred odor of each neuron was designated using a subset of the trials (n = 5 per odor), in the epoch under consideration (e.g., sample), based on the sign of the difference in the means, irrespective of statistical significance. These trials were then excluded and the odor selectivity was computed as the spike rate to preferred odor minus the nonpreferred odor on the remaining data in a sliding bin of 100 ms. All neurons from each of the three areas are included in Fig. 3A (left graph). In Fig. 4I-N, all neurons from each of the three areas are included.

For the decoding analysis (Fig. 3B-D, 4O-R and 6A, B), we trained a support vector machine (SVM) (Fan, Rong-En, 2008) with neural responses recorded simultaneously from Pir, OFC, or ALM to classify sample odor identity, trial type, or match/nonmatch. 48-120 neurons were simultaneously recorded from Pir, 16-82 neurons from OFC and 14-82 neurons from ALM. In Fig. 3B, we used the spike counts in a sliding bin of 500 ms from 40 randomly selected neurons from each of the three areas at 100 ms steps. Sessions with insufficient number of simultaneously recorded neurons were excluded. In Fig. 3C, we used a 500 ms sliding bin with neural responses from all Pir neurons recorded in a session. In Fig. 3D, we used the firing rates in the 500 ms time window before animal’s first lick. The decoding capability of each area was estimated by using varying numbers of randomly selected neurons that are recorded simultaneously in a session. As we included more neurons, sessions with insufficient number of neurons drop out in the decoding analysis. The classifier was trained on randomly selected 90% of the trials in each session and then tested on the remaining 10% of the trials. Only correct trials were used. The training/testing was repeated 50 times for every given number of neurons and for all the sessions that may be included. The correct rates from the 50 repetitions were then averaged. When comparing performance in control and inactivation conditions (Fig. 4O-R), the classifiers were trained on correct control trials and tested on correct laser trials and held-out correct control trials.

### Imaging data analysis

The raw images were first motion corrected with SIMA package (Kaifosh et al., 2014) (Release 1.3.2) and verified manually. Regions of interest (ROIs) were selected automatically with constrained nonnegative matrix factorization (CNMF) (Pnevmatikakis et al., 2016). The CNMF algorithm infers the time-varying background and extracts smoothed ΔF/F signals, which were used for plotting only. For data analysis, we manually computed unfiltered ΔF/F traces as follows. We obtained the raw fluorescent trace of each ROI by applying the spatial component (ROI filter) on the image sequence. We then smooth the raw trace in each trial with a 1-s averaging window (boxcar) and take the minimal fluorescence value in the inter-trial interval as the baseline. The ΔF/F signals were calculated by subtracting the baseline from the raw trace and dividing the difference by the baseline. We used the constrained deconvolution spike inference algorithm (FOOPSI) in the CNMF package to infer the spikes. The decay time constant was set at 0.7 second. The deconvolved activity was then smoothed using a gaussian filter over a five-element sliding window.

We compared the means of the response to odors A and B during the sample and delay epochs using a Mann-Whitney U test. Each trial contributed a scalar value: the average ΔF/F signal from t = 0-2.5 s from onset of the sample odor. Neurons were classified as sample odor-selective if *P* < 0.01 (two-tailed, not corrected for multiple comparisons). Test odor and choice selectivity were determined similarly using time bins of 2.5-4.0 s and 2.5-5.0 s from sample onset, respectively. Selectivity indices (SIs, Eq. 1) were computed from the same scalar values. The distributions of the sample odor SIs of ALM L2 neurons acquired with calcium imaging and those of ALM neurons sampled by electric recording were compared using a two-sample Kolmogorov-Smirnov test.

The standardized odor and choice selectivity (Fig. 5A and S8D) were computed by dividing the absolute value of the difference between the mean ΔF/F responses to odor A and B (or match and non-match) by the combined standard deviation. The combined standard deviation is the square root of the sum of the variances for the ΔF/F responses to odor A and B (or match and non-match).

To determine the duration of the calcium response, we first computed the mean and standard deviation of the baseline (the epoch before stimulus onset). Calcium transients were then identified as any response greater than 2 standard deviations away from the baseline and lasting for at least 100 ms. The peak time is the time of the maximum calcium response. Only the calcium transients during the sample and delay epochs were considered in Fig. 5E. We used identical methods to determine the duration and peak time of the inferred spiking activity except that a minimal response duration was not required.

To establish that a representation of sample odor in the population transient L2 neurons spans the entire sample plus delay, we measured the pairwise correlation in the peak times of the responses (Fig. S6A). Statistical significance of each r-value was established using Fisher z-transformed values and their s.e., without correction for multiple comparisons (gray shaded histograms, Fig. S6B). We also estimated the distribution of correlation coefficients expected under the null hypothesis, using a shuffle control. The calculations are identical except each ordered pair from the two neurons comprises peak times from non-corresponding trials. The red dashed distribution (Fig. S6B) was estimated using 1000 iterations of this procedure.

For the decoding analysis (Fig. 6A-B), we trained a support vector machine (SVM) (Fan, Rong-En, 2008) using the calcium responses to classify sample odor identity (Fig. 6A) or match/nonmatch (Fig. 6B). The calcium responses were computed as the average ΔF/F in a sliding bin of 100 ms at 100 ms steps using 40 randomly selected neurons from each of the five cortical depths in ALM.

We characterized a trial-by-trial association between the Ca responses of L2 neurons and the likelihood that the mouse would make an error. The signals themselves are indirect measurements of neural activity and highly skewed. We therefore applied a variant of the choice probability measure based on ordinal statistics. We included 109 neurons that had statistically reliable preference for sample odor A or B using the integrated Ca signal over the sample and delay epochs. We included both correct and error trials in determining the sample odor selectivity, to avoid biases favoring neurons with larger responses in correct trials due to random fluctuation. For each neuron, the integrated signal was ranked and scored as a percentile ranking on (0,1] using the trials in which the sample odor was the preferred odor to form the vector of trial-by-trial responses for each neuron, 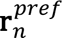, where the subscript identifies the neuron. The percentiles were assigned for each neuron independently (not the population). The percentiles from all the neurons were then pooled. We used the same procedure using the responses to each neuron’s nonpreferred odor to form 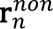.

The heatmap in Fig. 6E was formed by parametrically varying the two criteria, *κ_pref_* and *κ_non_*, to compute the proportion of errors when **r***^pref^* < *κ_pref_* and **r***^non^* > *k_non_*, using the combined data from all 109 neurons (i.e., concatenating across all *n*). To evaluated the null hypothesis that these responses have only a random association with the behavioral outcome, we conducted a simple logistic regression using the percentiles themselves:

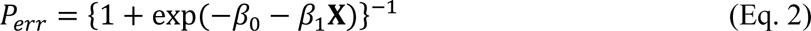

Where X is the vector of the transformed percentiles:

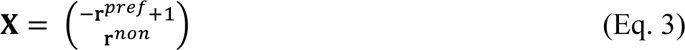

Note that the percentiles are simply reversed for **r***^pref^* so that the larger percentiles correspond to the weakest responses. We report the p-value associated with {*H_0_*: *β*_1_ = 0}.

### Behavior/inhibition data analysis

Mouse performance (P) was reported as the fraction of correct responses in all trials. Animals may perform below chance (50%) due to “no choice” trials (e.g., when they are challenged with novel C/D × C/D pairs; Fig. S1C). To assess the statistical reliability of photoinhibition on P we generated the distribution of log odds (𝓛) under the null hypothesis

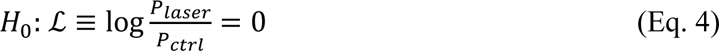

We randomly permuted the designations, correct/incorrect, among the laser and control trials within each trial type (AA/AB/BB/BA), repeating the process 10,000 times. From this distribution, we obtain the two-tailed probability of obtaining the observed log odds under *H*_0_.

To compare the effect of inactivation across tasks, we generated a distribution of 𝓛 for each task by bootstrapping. The laser and control trials were re-sampled respectively with replacement within each trial type (AA/AB/BB/BA). Repeating this process 10,000 times, we established a *t*-statistic from the means and variances of these distributions (degree of freedom based on the number of experimental trials). The reported significance reflects one-tailed comparisons.

We used the following logistic model to characterize an animal’s bias. “No choice” trials in which animals did not respond were excluded in this analysis (< 1% of trials).

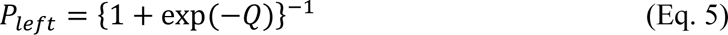

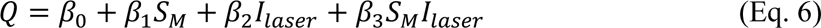

Where *P_left_* is the probably that the animal licks to the left port, *S_M_* is +1 or −1 if the trial is a match or non-match, respectively, and *I_L_* is 1 or 0 for laser on or off, respectively. The beta terms are fitted coefficients: *β*_0_ quantifies the bias in favor of left on control trials in units of log odds; *β*_0_ + *β*_2_ quantifies the bias in laser-on trials; *β*_1_ quantifies how well the animal uses the match/non-match information to determine the direction of licking (i.e., sensitivity to condition) on the control trials; and *β*_1_ + *β*_3_ quantifies the sensitivity in laser-on trials. In a well-trained animal, *β*_1_ is always positive. *β_2_* is an estimate of the side bias induced by inactivation and *β_3_* is an estimate of the impairment on sensitivity by inactivation.

When trials from all the animals are combined in this analysis, the bias coefficients could be underestimated (*β_0_*, *β_2_*) because the bias for left or right is different across experiments. It is theoretically possible that underestimation of the laser induced bias, *β_2_*, could lead to misattribution of this effect to a laser induced change in sensitivity, *β*_3_. To address this, we first determined the bias of each animal in the laser condition by comparing the rate of correct match and non-match trials. Based on this bias, we designated left or right as the *preferred* lick port and match or non-match as the preferred trial for each animal. Then we combined the trials from all the animals to fit the following modified model:

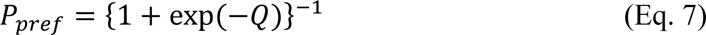

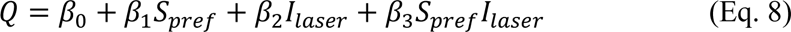

Where *P_pref_* is the probability that the animal licks to the preferred port, and *S_pref_* is +1 or −1 if the trial is preferred or non-preferred, respectively. In this model, *β*_2_ estimates the side bias induced by inactivation across all sessions. Importantly, if inactivation only biased the mouse to lick more to one port or the other—a different bias on each experiment—this procedure would fail to reject the null hypothesis, H_0_: *β*_3_ = 0. We complemented this analysis using Monte Carlo methods to estimate the magnitude of impairment on the task that is not accounted for by a bias to the left or right lick port. We simulated datasets using the estimated *β* coefficients and their standard error, while setting *β*_2_ = 0. This recovers the proportion correct on the control trials and models the proportion correct on the inactivation trials, were no bias induced by the laser. In Fig. S4C, we show the distribution of impairments from 10,000 repetitions.

### Attractor network models

We constructed recurrent neural network models consisting of three areas (Fig. 7A). Each area *ɑ* = 1,2,3 (representing Pir, intermediate areas, and ALM, respectively) contains *N* = 80 units whose activities are represented by an *N*-dimensional vector **x***_a_* and follow the dynamics

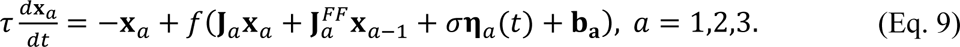

The matrices **J***_a_* and 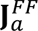 represent recurrent input and feedforward input from the previous area, respectively. The vector **x**_0_ is two-dimensional, and its two elements are indicator variables (with value 1 or 0) representing the presence or absence of odors A and B. The term *σ***η***_a_*(*t*) represents independent white noise input with standard deviation σ. The vector **b***_a_* represents the bias inputs to each unit. The function *ƒ* is rectified-linear and the time constant τ (which represents combined membrane and synaptic time constants) equals 100 ms.

In each trial, the network receives sample and test odors (A or B) for 500 ms each, beginning at times *t* = 1 and 3 s. Each trial is drawn randomly from one of the four trial types. A readout of the network must classify the trial as match or non-match during the response period, from *t* = 3.5 s to *t* = 4 s. The readout is a softmax function of the ALM activity **x**_3_. At the beginning of each trial, the initial values of the **x***_a_* vectors are taken to be independent rectified Gaussian random variables with standard deviation 0.05. Networks are simulated with a timestep of 20 ms.

The networks are trained through gradient descent with TensorFlow and the Adam optimizer (Agarwal et al., 2016; Kingma and Ba, 2014) to minimize a loss *L* = *L_classifier_* + *L_activity_* determined by the classifier and an activity regularization term. Specifically, *L_classifier_* in a given training epoch equals the summed cross-entropy loss between the classifier’s output and the desired output (match or non-match) during the response period, averaged over a batch size of 50 trials. The regularization term *L_activity_* = 10^−4^ · 〈||**r**_*a*_||^2^〉 is proportional to the *ℓ*_2_ norm of the activities averaged across units, time, areas, and batches. Every 50 epochs of training, the network is tested on 1000 trials to determine its classification accuracy, and training ceases when this accuracy exceeds 95%. The noise σ equals 0.05 during training and 0.1 during this testing phase. The learning rate of the optimizer decreases logarithmically from 10^−3^ to 10^−4^ over 1000 epochs. Networks which do not reach the 95% criterion accuracy after these 1000 epochs are discarded.

At the beginning of training, all recurrent and feedforward weights **J***_a_*, 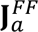 are initialized as independent random Gaussian variables with standard deviation 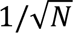 (except for 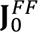, which has standard devation 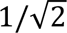 because of the dimension of **x**_0_). Half of the feedforward weights are then randomly set equal to zero, representing sparser connections across areas versus within areas. The softmax classifier weights are initialized with standard deviation 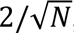, and the biases **b***_a_* are initialized to zero. We assume that only **J***_a_* and **b*_a_*** are learned. All other variables are fixed during training.

To generate the performance curves in Fig. 7B, 60 trained networks were tested with σ = 0.2 under four conditions. In the control condition, the dynamics were identical to those described above. In the other conditions, which mimic ALM inactivation during different periods of the task, an inhibitory input was applied to ALM units during the sample and early delay periods (the 1 s following sample onset), the late delay period (the final 1 s of the delay period), or the entire sample and delay periods. The inhibitory input was equivalent to reducing the biases of all ALM units **b**_3_ by 5. To generate the curve for networks without ALM persistence, modifications of **J***_a_* were restricted to only **J**_1_ and **J**_2_ while **J**_3_ was held fixed, so that the training algorithm could not learn to implement attractor dynamics in ALM (*a* = 3). Conversely, for networks with only ALM persistence, training was restricted to only **J**_3_, while **J**_1_ and **J**_2_ were held fixed.

## References

Amit, D.J., and Brunel, N. (1997). Model of global spontaneous activity and local structured activity during delay periods in the cerebral cortex. Cereb. Cortex 7, 237–252.

Bechara, A., Damasio, H., and Damasio, A.R. (2000). Emotion, Decision Making and the Orbitofrontal Cortex. Cereb. Cortex 10, 295–307.

Bittner, K.C., Milstein, A.D., Grienberger, C., Romani, S., and Magee, J.C. (2017). Behavioral time scale synaptic plasticity underlies CA1 place fields. Science (80-.). 357, 1033–1036.

Carandini, M., and Churchland, A.K. (2013). Probing perceptual decisions in rodents. Nat. Neurosci. 16, 824–831.

Chen, T.W., Wardill, T.J., Sun, Y., Pulver, S.R., Renninger, S.L., Baohan, A., Schreiter, E.R., Kerr, R.A., Orger, M.B., Jayaraman, V., et al. (2013). Ultrasensitive fluorescent proteins for imaging neuronal activity. Nature 499, 295–300.

Chen, T.W., Li, N., Daie, K., and Svoboda, K. (2017). A Map of Anticipatory Activity in Mouse Motor Cortex. Neuron 94, 866–879.e4.

Cisek, P. (2011). Cortical mechanisms of action selection: the affordance competition hypothesis. Model. Nat. Action Sel. 208–238.

Cisek, P., and Kalaska, J.F. (2005). Neural correlates of reaching decisions in dorsal premotor cortex: Specification of multiple direction choices and final selection of action. Neuron 45, 801–814.

Evarts, E. V., and Tanji, J. (1976). Reflex and intended responses in motor cortex pyramidal tract neurons of monkey. J. Neurophysiol. 39, 1069–1080.

Freedman, D.J., and Assad, J.A. (2006). Experience-dependent representation of visual categories in parietal cortex. Nature 443, 85–88.

Funahashi, S., Bruce, C.J., and Goldman-Rakic, P.S. (1989). Mnemonic coding of visual space in the monkey’s dorsolateral prefrontal cortex. J. Neurophysiol. 61, 331–349.

Fuster, J.M. (1973). Unit activity in prefrontal cortex during delayed-response performance: neuronal correlates of transient memory. J. Neurophysiol. 36, 61–78.

Fuster, J.M., and Alexander, G.E. (1971). Neuron activity related to short-term memory. Science 173, 652–654.

Giessel, A.J., and Datta, S.R. (2014). Olfactory maps, circuits and computations. Curr. Opin. Neurobiol. 24, 120–132.

Gold, J.I., and Shadlen, M.N. (2003). The influence of behavioral context on the representation of a perceptual decision in developing oculomotor commands. J. Neurosci. 23, 632–651.

Gold, J.I., and Shadlen, M.N. (2007). The Neural Basis of Decision Making. Annu. Rev. Neurosci. 30, 535–574.

Goldman-Rakic, P.S. (1995). Cellular basis of working memory. Neuron 14, 477–485.

Guo, Z., Li, N., Huber, D., Ophir, E., Gutnisky, D., Ting, J., Feng, G., and Svoboda, K. (2014). Flow of cortical activity underlying a tactile decision in mice. Neuron 81, 179–194.

Guo, Z. V., Inagaki, H.K., Daie, K., Druckmann, S., Gerfen, C.R., and Svoboda, K. (2017). Maintenance of persistent activity in a frontal thalamocortical loop. Nature.

Hanks, T.D., Kopec, C.D., Brunton, B.W., Duan, C.A., Erlich, J.C., and Brody, C.D. (2015). Distinct relationships of parietal and prefrontal cortices to evidence accumulation. Nature 520, 220–223.

Harvey, C., Coen, P., and Tank, D. (2012). Choice-specific sequences in parietal cortex during a virtual-navigation decision task. Nature.

Jones, E.G. (1998). Commentary Viewpoint : the Core and Matrix of Thalamic. Neuroscience 85, 331–345.

Kadohisa, M., Petrov, P., Stokes, M., Sigala, N., Buckley, M., Gaffan, D., Kusunoki, M., and Duncan, J. (2013). Dynamic Construction of a Coherent Attentional State in a Prefrontal Cell Population. Neuron 80, 235–246.

Larkum, M.E., Zhu, J.J., and Sakmann, B. (1999). A new cellular mechanism for coupling inputs arriving at different cortical layers. Nature 398, 338–341.

Larkum, M.E., Nevian, T., Sandler, M., Polsky, A., and Schiller, J. (2009). Synaptic integration in tuft dendrites of layer 5 pyramidal neurons: a new unifying principle. Science 325, 756–760.

Li, N., Chen, T.-W., Guo, Z. V., Gerfen, C.R., and Svoboda, K. (2015). A motor cortex circuit for motor planning and movement. Nature 519, 51–56.

Liu, D., Gu, X., Zhu, J., Zhang, X., Han, Z., Yan, W., Cheng, Q., Hao, J., Fan, H., Hou, R., et al. (2014). Medial prefrontal activity during delay period contributes to learning of a working memory task. Science (80-.). 346, 458–463.

Luo, L., Callaway, E.M., and Svoboda, K. (2018). Genetic Dissection of Neural Circuits: A Decade of Progress. Neuron 98, 256–281.

Madisen, L., Garner, A.R., Carandini, M., Zeng, H., Madisen, L., Garner, A.R., Shimaoka, D., Chuong, A.S., Klapoetke, N.C., Li, L., et al. (2015). Transgenic Mice for Intersectional Targeting of Neural Sensors and Effectors with High Specificity and Performance NeuroResource Transgenic Mice for Intersectional Targeting of Neural Sensors and Effectors with High Specificity and Perf. 942–958.

Mainen, Z.F., and Kepecs, A. (2009). Neural representation of behavioral outcomes in the orbitofrontal cortex. Curr. Opin. Neurobiol. 19, 84–91.

Mongillo, G., Barak, O., and Tsodyks, M. (2008). Synaptic Theory of Working Memory. Science (80-.). 319, 1543–1546.

Padoa-Schioppa, C., and Assad, J. a (2006). Neurons in the orbitofrontal cortex encode economic value. Nature 441, 223–226.

Polsky, A., Mel, B.W., and Schiller, J. (2004). Computational subunits in thin dendrites of pyramidal cells. Nat. Neurosci. 7, 621–627.

Ramus, S.J., and Eichenbaum, H. (2000). Neural correlates of olfactory recognition memory in the rat orbitofrontal cortex. J. Neurosci. 20, 8199–8208.

Romo, R., Brody, C.D., Hernández, a, and Lemus, L. (1999). Neuronal correlates of parametric working memory in the prefrontal cortex. Nature 399, 470–473.

Selen, L.P.J., Shadlen, M.N., and Wolpert, D.M. (2012). Deliberation in the Motor System: Reflex Gains Track Evolving Evidence Leading to a Decision. J. Neurosci. 32, 2276–2286.

Shadlen, M.N., and Kiani, R. (2013). Decision making as a window on cognition. Neuron 80, 791–806.

Shadlen, M.N., and Newsome, W.T. (2001). Neural Basis of a Perceptual Decision in the Parietal Cortex (Area LIP) of the Rhesus Monkey. J. Neurophysiol. 86, 1916–1936.

Shadlen, M.N., and Shohamy, D. (2016). Decision Making and Sequential Sampling from Memory. Neuron 90, 927–939.

Snyder, L.H., Batista, A.P., and Andersen, R.A. (1997). Coding of intention in the posterior parietal cortex. Nature 386, 167–170.

Sosulski, D.L., Bloom, M.L., Cutforth, T., Axel, R., and Datta, S.R. (2011). Distinct representations of olfactory information in different cortical centres. Nature 472, 213–216.

Spivey, M.J., Grosjean, M., and Knoblich, G. (2005). From The Cover: Continuous attraction toward phonological competitors. Proc. Natl. Acad. Sci. 102, 10393–10398.

Svoboda, K., and Li, N. (2018). Neural mechanisms of movement planning: motor cortex and beyond. Curr. Opin. Neurobiol. 49, 33–41.

Yang, T., and Shadlen, M.N. (2007). Probabilistic reasoning by neurons. Nature 447, 1075–1080.

Yang, G.R., Murray, J.D., and Wang, X.-J. (2016). A dendritic disinhibitory circuit mechanism for pathway-specific gating. Nat. Commun. 7, 12815.

Zhao, S., Ting, J.T., Atallah, H.E., Qiu, L., Tan, J., Gloss, B., Augustine, G.J., Deisseroth, K., Luo, M., Graybiel, A.M., et al. (2011). Cell type–specific channelrhodopsin-2 transgenic mice for optogenetic dissection of neural circuitry function. Nat. Methods 8, 745–752.

Zingg, B., Hintiryan, H., Gou, L., Song, M.Y., Bay, M., Bienkowski, M.S., Foster, N.N., Yamashita, S., Bowman, I., Toga, A.W., et al. (2014). Neural networks of the mouse neocortex. Cell 156, 1096–1111.

## Methods references

Agarwal, A., Barham, P., Brevdo, E., Chen, Z., Citro, C., Corrado, G.S., Davis, A., Dean, J., Devin, M., Ghemawat, S., et al. (2016). TensorFlow: Large-Scale Machine Learning on Heterogeneous Distributed Systems. ArXiv 1603.04467, [cs].

Fan, Rong-En, et al (2008). LIBLINEAR: A library for large linear classification. J. Mach. Learn. Res. 9, 1871–1874.

Guo, Z., Li, N., Huber, D., Ophir, E., Gutnisky, D., Ting, J., Feng, G., and Svoboda, K. (2014a). Flow of cortical activity underlying a tactile decision in mice. Neuron 81, 179–194.

Guo, Z. V, Hires, S.A., Li, N., O’Connor, D.H., Komiyama, T., Ophir, E., Huber, D., Bonardi, C., Morandell, K., Gutnisky, D., et al. (2014b). Procedures for behavioral experiments in head-fixed mice. PLoS One 9, e88678.

Kaifosh, P., Zaremba, J.D., Danielson, N.B., and Losonczy, A. (2014). SIMA: Python software for analysis of dynamic fluorescence imaging data. Front. Neuroinform. 8, 80.

Kingma, D.P., and Ba, J.L. (2014). Adam: A Method for Stochastic Optimization. ArXiv 1412.6980, [cs].

Pachitariu, M., Steinmetz, N.A., Kadir, S.N., Carandini, M., and Harris, K.D. (2016). Fast and accurate spike sorting of high-channel count probes with KiloSort. NIPS Proc. 4448–4456.

Pnevmatikakis, E.A., Soudry, D., Gao, Y., Machado, T.A., Merel, J., Pfau, D., Reardon, T., Mu, Y., Lacefield, C., Yang, W., et al. (2016). Simultaneous Denoising, Deconvolution, and Demixing of Calcium Imaging Data. Neuron 89, 299.

Root, C.M., Denny, C.A., Hen, R., and Axel, R. (2014). The participation of cortical amygdala in innate, odour-driven behaviour. Nature 515, 269–273.

Rossant, C., Kadir, S.N., Goodman, D.F.M., Schulman, J., Hunter, M.L.D., Saleem, A.B., Grosmark, A., Belluscio, M., Denfield, G.H., Ecker, A.S., et al. (2016). Spike sorting for large, dense electrode arrays. Nat. Neurosci. 19, 634–641.

